# Genetically recoding respiratory syncytial virus to visualize nucleoprotein dynamics and virion assembly

**DOI:** 10.1101/2024.05.12.593692

**Authors:** Margaret D. Mitrovich, Michael D. Vahey

**Affiliations:** Department of Biomedical Engineering, Washington University in St. Louis, St. Louis, Missouri, USA; Center for Biomolecular Condensates, Washington University in St. Louis, St. Louis, Missouri, USA

## Abstract

RNA viruses possess small genomes encoding a limited repertoire of essential and often multifunctional proteins. Although genetically tagging viral proteins provides a powerful tool for dissecting mechanisms of viral replication and infection, it remains a challenge. Here, we leverage genetic code expansion to develop a recoded strain of respiratory syncytial virus (RSV) in which the multifunctional nucleoprotein is site-specifically modified with an unnatural amino acid. The resulting virus replicates exclusively in cells capable of amber stop codon suppression and is amenable to labeling with tetrazine-modified fluorophores, achieving high signal-to-background. We use this tool to visualize the transfer of nucleoprotein complexes from cytoplasmic condensates directly to budding viral filaments at the cell surface and to cytoplasmic compartments containing viral surface proteins, suggesting multiple pathways for viral assembly.

---

Techniques to install versatile, endogenous tags into viral genomes would advance understanding of the spatiotemporal dynamics of viral replication and spread. While fluorescent fusion proteins and self-labeling tags including Halo, SNAP, and CLIP^1,2^ have become widely adopted in cell biology, they remain comparatively rare in virology research due to the technical challenges of successfully installing these relatively large (∼20-30 kDa) tags endogenously within viral genomes. Even when robust reverse genetics systems are available – as is the case for many viruses^3–6^ - the challenge remains of installing the tag without disrupting the structure and function of the protein of interest. This is particularly challenging in RNA virus research, where compact genomes (typically 10-30kbp in size) require that nearly every protein is essential and frequently performs diverse functions throughout the replication cycle. Additionally, genomic RNAs frequently play structural roles during virus infection and assembly^7–9^, further constraining the modifications that can be tolerated.

Genetic code expansion (GCE) offers several advantages that could help to overcome these challenges^10^. GCE enables the site-specific installation of a non-canonical amino acid (ncAA) through reassignment of the Amber stop codon. The resulting tag can have diverse biochemical properties, and is comprised of a single residue, minimizing its footprint and potential disruption to protein or RNA function. This approach has been successfully used in mammalian cells to mimic post-translational modifications^11^, to fluorescently label cellular proteins^12^, to map protein interaction interfaces^13^, and to develop covalent protein drugs^14^. Efforts to leverage GCE for the endogenous modification of proteins within the genome of a replicative virus have focused primarily on controlling replication for vaccine applications^15–18^, and recently GCE has been utilized to monitor HIV entry pathways^19^. However, using GCE to directly image virus replication and assembly within infected cells remains a challenge.

We sought to establish a conditionally-replicative strain of respiratory syncytial virus (RSV) with a genetically recoded nucleoprotein (N). RSV imposes a continued burden on human health and its mechanisms of assembly and entry remain unresolved^20–22^. N does not tolerate GFP tags on either its N- or C-termini^23^, and we found that even small peptide tags like the five-residue tetra-cysteine motif^24^ are lethal to RSV replication. We reasoned that GCE would avoid the disruption caused by even small insertions within N if we could overcome some of its challenges. RSV N is expressed at high levels beginning early in infection, and it is an essential co-factor for the transcription of RSV genes and replication of the RSV genome. Inefficient amber suppression is therefore likely to be lethal to viral replication and must be avoided. Using an optimized engineered pyrrolysyl tRNA/AARS pair^25^, we sought to determine sites in RSV N that lead to efficient amber suppression. To identify candidate residues, we used iPASS^26^and further refined results based on their surface accessibility within the structure of the assembled nucleocapsid^27,28^ (Figure 1A). Based on these criteria, we selected nine sites for testing using a BFP mini-genome: K7, L12, H43, F61, Y135, L139, D173, I183, and T340. For leading candidates identified by iPASS, we introduced silent mutations two residues upstream and downstream from the amber stop codon to enhance predicted amber suppression. Variants D175*, I183*, and T340* showed the highest transcriptional activity based on BFP expression, reaching levels several-fold higher than wildtype N when using Lys(Z) and comparable to wildtype when using TCO-K (Figure 1B).

**Figure 1:**
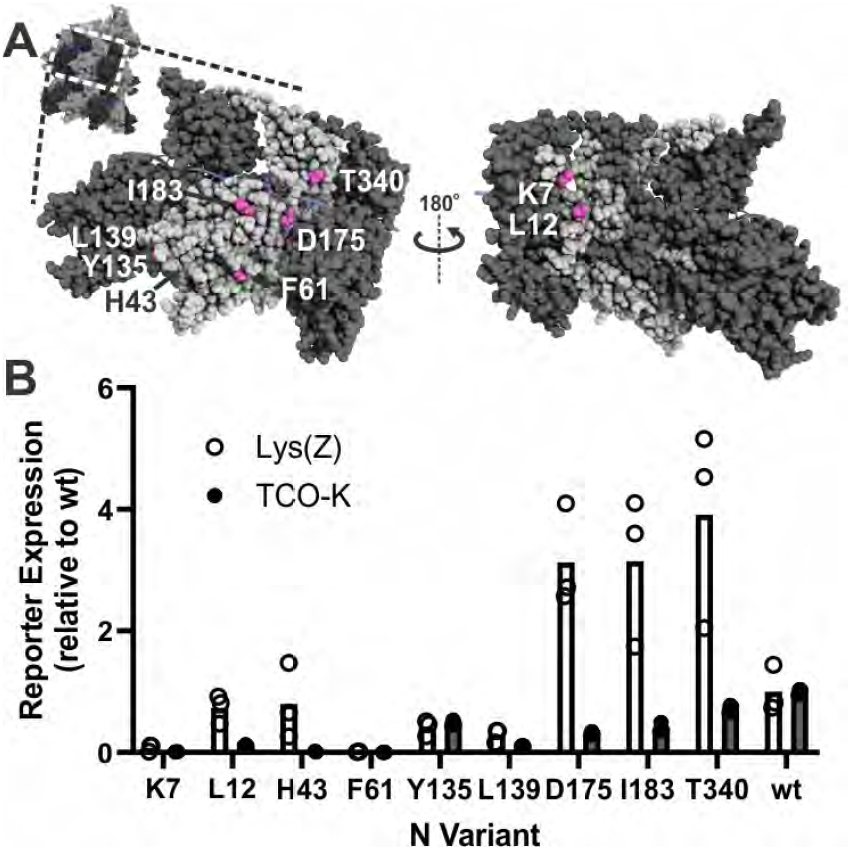
Identifying permissive sites for amber suppression in RSV N. (A) Structure of the RSV nucleocapsid (from PDB ID 4BKK), highlighting candidate sites for amber suppression (magenta). Candidate sites include residues exposed on both the outer and inner surfaces of the helical nucleocapsid. (B) Transcriptional activity of genetically recoded N relative to wildtype N, measured via expression of BFP from a mini viral genome.

We proceeded to rescue viruses possessing the two top-performing constructs from the mini-replicon assay, I183* and T340*. We inserted I183* and T340* mutations into a BAC containing the RSV genome with additional tags on the F and G proteins as well as a BFP reporter^29^ (Figure 2A; complete sequence available in Supporting Information) and transfected this along with helper plasmids into a stable BHK-21 line expressing the PylT/RS from a piggyBac-based cassette (‘BHK-M15’; complete sequence available in Supporting Information). While expression of the BFP reporter in cells transfected with the T340* variant remained low after several passages, the I183* variant showed robust replication. Although the rescued virus was able to infect and replicate in BHK-M15 cells, wildtype BHK-21 cells infected with equal amounts of virus showed no BFP expression at 48h post-infection (Figure 2B). This demonstrates that RSV strains possessing the I183* mutation in N are conditionally-replicative.

**Figure 2:**
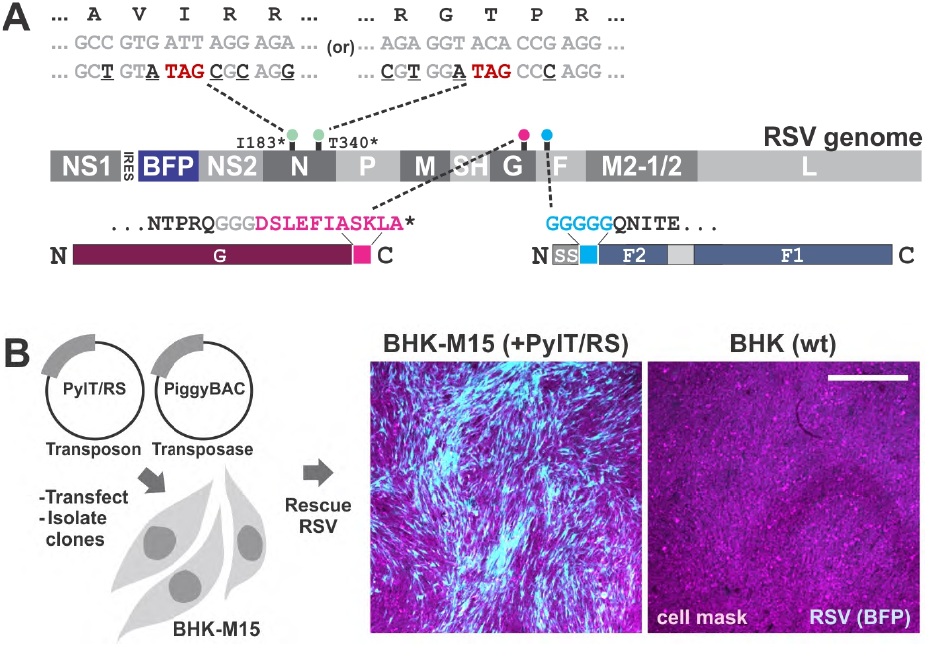
A genetically-recoded RSV strain. (A) Organization of the engineered RSV genome, including the BFP reporter expressed from an internal ribosome entry site, I183 / T340 amber mutations within N (green), and tags for enzymatic labeling on G and F (magenta and cyan). Engineered BACs contain either the N(I183*) or N(T340*) mutations. Underlined nucleotides adjacent to amber mutations in N indicate silent mutations predicted to enhance amber suppression. (B) Construction of BHK-M15 cells and rescue of RSV N(I183*). Only cells expressing the machinery for amber suppression support replication of recoded RSV, indicated by expression of the BFP reporter. Images are collected at 48h post-infection with RSV N(I183*) at matching multiplicity of infection and displayed at matching contrast levels. Scale bar = 1mm.

Although stable delivery of the PylT/RS cassette using a piggyBac transposon facilitates virus rescue, it limits infection studies to immortalized cell lines that tolerate clonal expansion. To circumvent this constraint, we developed an AAV vector to transiently deliver the machinery for genetic code expansion (Figure 3A; complete sequence available in Supporting Information). AAV vectors can transduce diverse cells and tissues in vitro and in vivo, and have previously been used to expand the genetic code in the mouse brain^30^. To leverage these advantages, we created AAV vectors containing a single copy of the *Mm*PylRS (under the control of a CMV promoter) and three copies of the M15 PylT (under the control of U6 promoters). To test AAV vectors for amber suppression, we measured expression of BFP with a premature amber stop codon (H10*) following transduction with AAV1-serotyped vector under different conditions: in the presence or absence of different ncAAs (TCO-K and LysZ, both at 500 μM) as well as with or without the proteosome inhibitor MG-132, known to increase AAV transduction^31^ (Figure 3B). Consistent with our observations in the mini-replicon system, we observed the highest levels of BFP expression in cells supplemented with Lys(Z). Treatment with MG-132 also had a pronounced effect on AAV transduction, suggesting an additional means to increase amber suppression efficiency in hard-to-infect cells. Consistent with results in BHK-M15 cells, Vero E6 cells infected with AAV (day 0) were susceptible to infection with RSV N(I183*) at 2d post-AAV infection, whereas mock-infected cells were not (Figure 3C).

**Figure 3:**
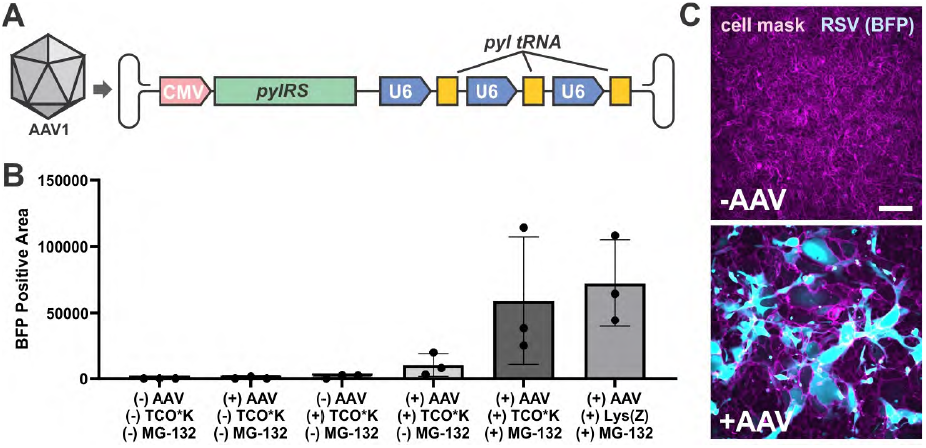
An AAV vector for genetic code expansion. (A) Schematic of the AAV1-serotyped vector. (B) Quantification of BFP signal from Vero-BFP(H10*) cells infected with AAV-M15 PylT/RS in the presence of the indicated supplements. (C) Images of cells infected with RSV N(I183*), with or without prior infection with AAV. Scale bar = 100μm. Cells are imaged at 48h post-infection with RSV (4d post-infection with AAV). Channels are displayed at matching contrast levels.

We proceeded to test labeling of cells infected with RSV N(I183*) using tetrazine-modified derivatives of JF549, JF646, and SiR^32,33^. Infected cells were amenable to labeling by all three dyes, with budding virions and cytoplasmic condensates termed inclusion bodies (IBs)^34^ clearly visible (Figure 4A, Supplementary Movie S1). Labeling via tetrazine dyes overlapped with N-specific antibodies, but showed better penetrance into the interior of IBs (Figure 4B). Interestingly, the three dyes exhibited different preferences for labeling different RSV-associated structures. Compared to JF646 and SiR, JF549 exhibited greater labeling of virions (identified through colocalization with F) relative to inclusion bodies and cytoplasmic background (Figure S1).

**Figure 4:**
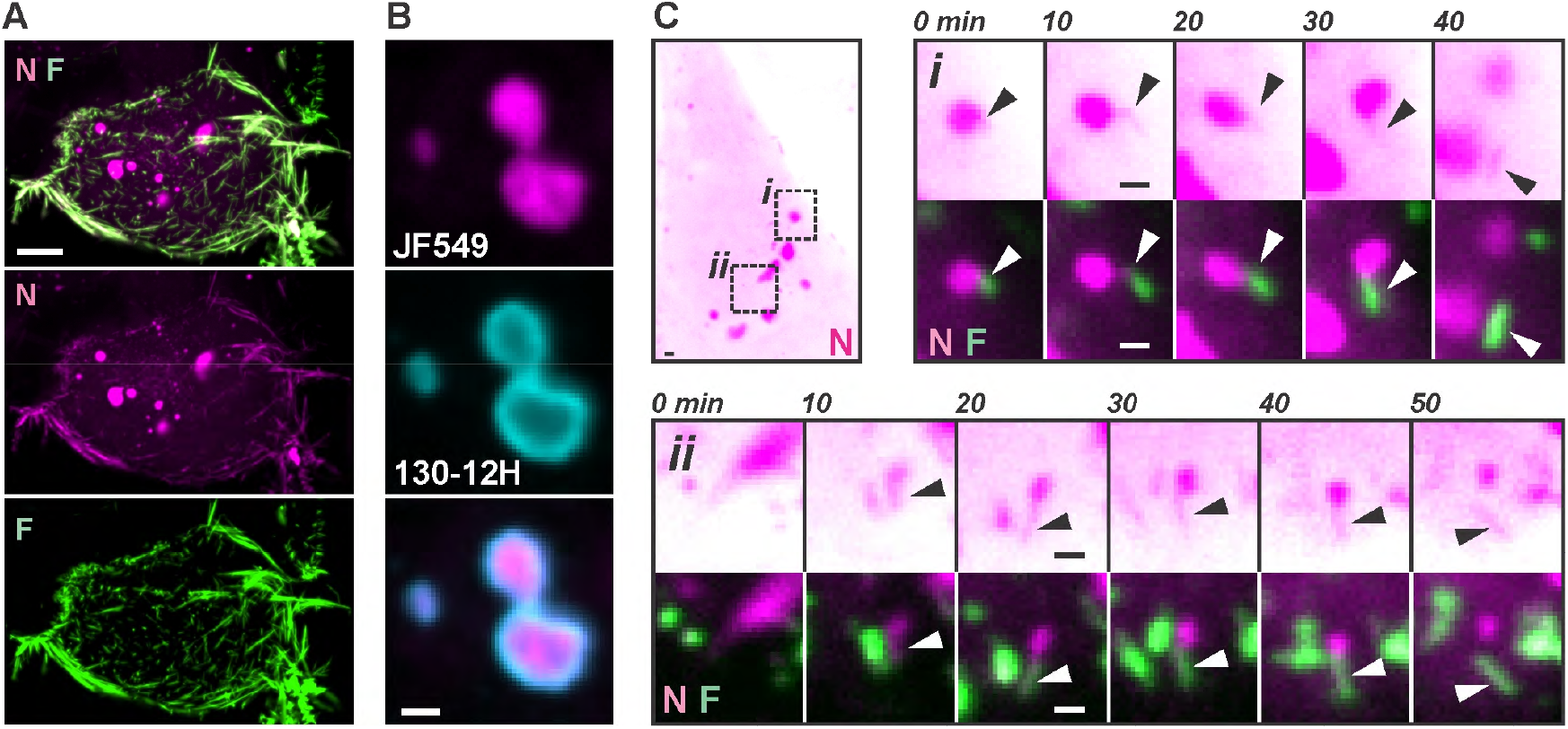
Nucleoprotein-containing structures emerge from dynamic inclusion bodies. (A) Image of an infected cell at 48h post-infection with RSV N(I183*), labeled with JF549-Tz (nucleoprotein) and a D25 Fab fragment (fusion protein). (B) Images of RSV inclusion bodies labeled with JF549-Tz or the monoclonal antibody 130-12H. (C) Time series of N filaments extending from inclusion bodies and associating with F at the cell surface. Scale bar = 1μm in all panels.

In addition to the engineered amber stop codon in N, the A2 strain of RSV contains natural amber stop codons in the membrane proteins SH and G. Since the C-termini of SH and G are both extracellular, we reasoned that amber suppression during translation of either protein would lead to detectable signal on the cell surface after labeling with membrane-impermeable dyes. However, cells labeled with membrane-impermeable sulfo-Cy3 tetrazine for 30 minutes showed only background labeling, in contrast to cells labeled with membrane-permeable JF549-Tz and imaged under the same conditions (Figure S2), suggesting that natural amber stop codons in SH and G are not suppressed to a level that is detectable via cell-surface fluorescence labeling.

Using time lapse microscopy, we observed the dynamics of IBs within infected cells. Consistent with previous reports, we observe both IB coalescence^23^ (Supplementary Movie S2) as well as the rapid emergence of small puncta and thread-like protrusions from larger inclusion bodies (Figure S3, Supplementary Movie S3)^35^. Using a high-affinity Fab fragment (D25) to visualize RSV F over time, we observed instances where thin protrusions began extending from IBs and became partially coated with F from their opposite end (Figure 4C).

These F-coated protrusions grew larger over time before separating from the IB. Interestingly, the maximal length of the N protrusion is ∼1.5μm, similar to the length of the full-length encapsidated genome predicted from the cryo-EM structure^27^, suggesting that these events may represent packaging of individual genomes from cytoplasmic condensates directly into budding virions.

In addition to co-localization of N with F at the plasma membrane, we also observed intracellular compartments that are positive for the D25 Fab and that peripherally associate with N puncta (Figure 5A). These compartments appear after prolonged incubation with D25, suggesting that they are comprised of F that has been recycled from the cell surface where the Fab fragment could bind. Because the D25 Fab exhibits exceptionally slow dissociation (*k*_*off*_ < 10^−5^ s^-1^)^36^, we expect it to bind stably to F throughout the duration of these experiments. The structures we observe resemble the assembly granules reported by others^20,22,37^. Using timelapse microscopy, we observed frequent interactions between inclusion bodies and these compartments (Supplementary Movie S4), as well as their acquisition of N-positive puncta during a glancing interaction with a larger inclusion body (Figure 5B).

**Figure 5:**
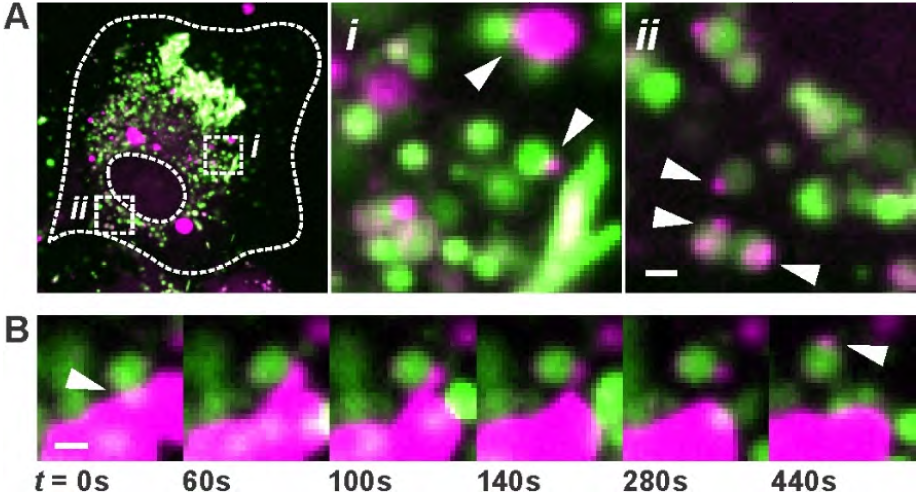
Nucleoprotein-containing foci associated with intracellular compartments containing F. (A) Vero cell infected with AAV at 0h, infected with RSV N(I183*) at 48h, and labeled and imaged at 84h. Insets *i* and *ii* show intracellular compartments labeled with the D25 Fab with nucleoprotein foci peripherally attached. (B) Time series showing the exchange of N between a large inclusion body and an F-containing compartment. Scale bar = 1μm in all panels.

This work establishes the feasibility of using genetic code expansion to image viral replication and assembly. Using this approach, we were able to observe packaging of viral nucleoprotein complexes into budding filaments, as well as their dynamic association with intracellular compartments containing the fusion protein F. The framework we present here is enabled by recent parallel advances to enhance the efficiency of amber suppression^25,26^ and the development of bright and photostable dyes for labeling intracellular proteins^32,33,38^. Live fluorescence imaging using genetic code expansion can suffer from high background resulting from labeling of residual charged tRNAs or from cellular proteins whose native amber stop codons are suppressed. In this work, these challenges are mitigated by the high local concentration of N within budding virions and inclusion bodies.

These features are conserved for nucleoproteins in the order *Mononegavirales*^39^, suggesting that the approach described here could be broadly applicable. The ability to deliver suitable tRNA/tRNA synthetase pairs stably (via transposase-mediated integration) or transiently (via AAV vectors) will facilitate initial virus rescue and support subsequent experiments using the conditionally replicative virus in vitro and potentially in vivo.

## Methods

### Identification of efficient sites for Amber suppression

We identified potential sites for amber stop codon insertion in N through three criteria: (1) amber suppression efficiency predicted by iPass^26^; (2) surface exposure of the residue within the assembled nucleocapsid^27^; and (3) biochemical similarity of the substituted residue to TCO-K. Nine sites were selected for testing and cloned into a codon-optimized N. We then transfected HEK293T cells with RSV minireplicon components (P, M2-1, L, T7 RNA Pol, and mutant N variant), M15 pylT/RS, and pT7 RSV-BFP at ratios of (2:2:2:1:2):2:4 respectively for a total of 200 ng of DNA per well for a 96 well plate. Cells were maintained in media containing either Lys(Z) or TCO-K at a concentration of 500μM. 24 hours post-transfection, cells were imaged and the expression of BFP was quantified.

### RSV reverse genetics and rescue

To support multi-cycle replication of RSV strains with amber stop codons, we constructed a cell line derived from BHK-21 using piggyBac transposase and a transposon vector containing the pyrrolysine tRNA synthetase from *Methanosarcina mazei* with mutations to accommodate TCO-K (Y30A, Y384F), puromycin resistance expressed from an IRES, and three copies of an optimized tRNA (‘M15’)^25^, following the general approach of Elsässer et al 2016^11^. After transfecting cells in a 6-well plate with Lipofectamine 2000, 2.5μg transposon vector, and 0.25 μg transposase vector, puromycin resistant cells were clonally expanded and tested for amber suppression by transfecting a plasmid encoding BFP with an amber stop codon substituting for His10. Clones showing high activity (‘BHK-M15’) were expanded and banked.

For the generation of recombinant RSV harboring specific mutations, we cloned I183* and T340* mutations into an RSV BAC using recombineering^4,40^. Mutations to BACs were verified by full-plasmid sequencing. BACs were transfected along with codon-optimized helper plasmids and T7 RNAP into BHK-M15 cells as previously described^4,29^, and cells were cultured in OptiMEM supplemented with 2% fetal bovine serum, antibiotic / antimycotic, and 100μM TCO-K. Cells were monitored for BFP expression, and expanded until ∼100% of cells were BFP-positive. Virus was collected by harvesting infected cells in PBS, subjecting them to a freeze-thaw cycle in liquid nitrogen, and pelleting cell debris at 1000×g. Viral stocks were flash frozen in liquid nitrogen and stored at -80°C until use.

### AAV production and transduction

Between 4-6 T75 flasks of HEK293T cells were transfected at 70-80% confluency with Helper, Rep/Cap (AAV1 serotype), and transfer vector (5 μg of each plasmid per T75 flask). Media was exchanged 24h post-transfection and AAV was harvested and purified from both culture media and transfected cells at 5d post-transfection using PEG/chloroform extraction^41^. Final AAV stocks were concentrated in OptiMEM and stored at 4°C until use. VeroE6 cells for experiments were infected with AAV overnight at 37°C in serum-free media. For quantification of AAV transduction and amber suppression, Vero E6 cells stably expressing BFP with an amber stop codon substituting for His10 were infected in the same manner and cultured with the supplements specified in Figure 3. Cells were imaged at 2 days post infection. Although proteasome inhibitors MG-132 and carfilzomib dramatically enhanced AAV transduction, we did not use these additives for cells to be used for RSV infection.

### RSV N(I183*) infection, labeling, and imaging

Vero E6 cells infected with AAV-M15 were maintained in 100μM TCO-K starting at 2d post-AAV-infection with daily exchanges of media to minimize loss of TCO reactivity over time. On day 2 or day 3 post-AAV-infection, cells were infected with frozen stocks of RSV N(I183*) by centrifugation at 1200×g for 10 minutes and subsequent incubation at 37°C for 4h before exchanging to fresh virus growth media with TCO-K.

Vero E6 cells infected with AAV and RSV were cultured in 8-chambered coverglass (Cellvis) for labeling and imaging. Beginning at 2-3h prior to labeling with tetrazine dyes, virus growth media containing 100μM TCO-K was replaced with fresh OptiMEM containing 0.5mM Lys(Z) and exchanged periodically. Immediately prior to labeling, fresh tetrazine dyes were diluted to 2μM in OptiMEM and added to cells for 30 minutes at 37° C. Residual dye was washed 3× with virus growth media supplemented with Lys(Z). For experiments using the D25 Fab to visualize RSV F, labeled Fab was added to virus growth media at a concentration of 2 nM and left present for the duration of imaging experiments to enable labeling of newly-synthesized F as it reaches the plasma membrane. Imaging was performed on a Nikon Ti2 microscope equipped with a CSU-X spinning disk, Hamamatsu Flash 4.0 camera, and 405/488/561/640 nm laser lines. Images were collected using either a 40x 1.30 NA or 60x 1.45 NA oil objective (Nikon). Cells were maintained at 37°C and 5% CO_2_ using a stage-top incubator (Tokai Hit) and objective heater. Images were analyzed using custom Matlab scripts and Fiji.

## Notes

### Competing Interest Statement

The authors have declared no competing interest.

