## Supporting Information for "Genetically recoding respiratory syncytial virus to visualize nucleoprotein dynamics and virion assembly"

**Supporting Sequences** .......... 6-11

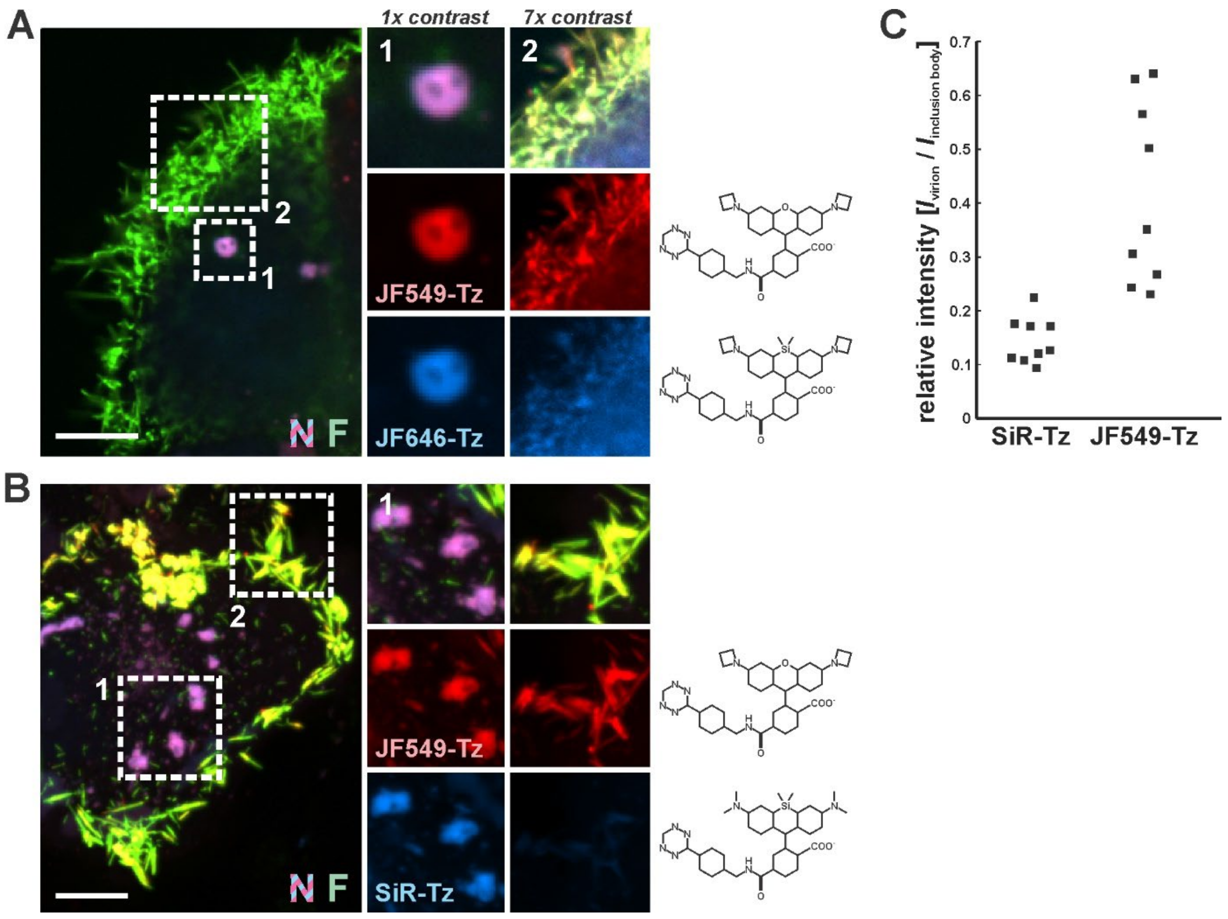

**Figure S1: Site-specific fluorescent labeling of RSV N in live cells.** (A) Confocal slice of a Vero cell infected with AAV-M15 (0h) and RSV (48h), and labeled simultaneously using JF646-Tz and JF549-Tz. Inset 1 shows an inclusion body and inset 2 shows virions budding at the cell surface. (B) Same as *A*, but with SiR-Tz in place of JF646-Tz. (C) Quantification of relative intensities for N labeled with JF549-Tz and SiR-Tz, as in panel *A*. Each data point represents average intensities for virions and IBs associated with a single cell. All scale bars = 10 µm.

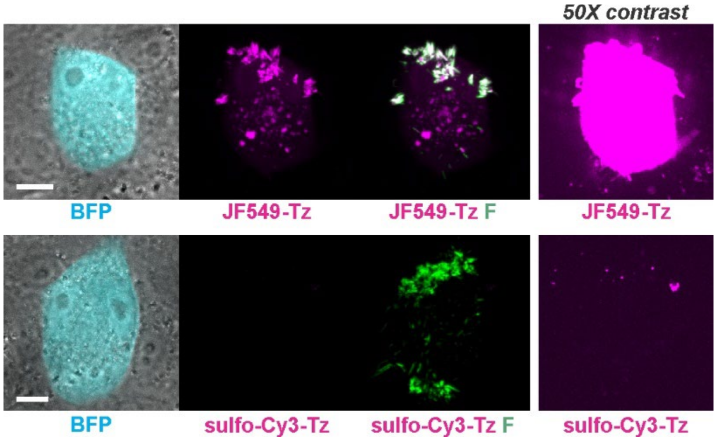

**Figure S2: Membrane-impermeable dyes do not label RSV N(I183*)-infected cells.** Cells infected with RSV N(I183*) are labeled with membrane permeable JF549-Tz (top row) or membrane-impermeable sulfo-Cy3-Tz (bottom row). Images are acquired using the same settings and displayed at the same contrast levels. Images to the right show the tetrazine channel with contrast enhanced 50-fold over the images to the left. Images are maximum-intensity projections. Scale bar = 10 μm.

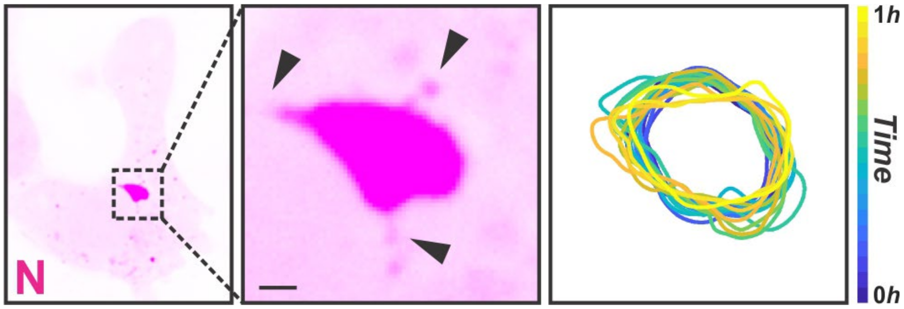

**Figure S3: Extrusion of N from RSV inclusion bodies.** Vero cell infected with AAV at 0h, infected with RSV N(I183*) at 48h, and labeled and imaged at 84h. Inset shows thin projections emerging from the surface of an inclusion body. Right: plot showing the boundaries of the same inclusion body to the left over the course of 1h. Scale bar = 1 μm.

**Supplementary Movie S1**: Fluorescence microscopy time series of a Vero cell infected with RSV N(I183*), labeled with JF549-Tz (N; magenta) and Alexa488-D25 Fab (F; green), and imaged at 10-minute intervals over the course of 250 minutes. Images are displayed as a maximum intensity projection. Scale bar = 10 μm.

**Supplementary Movie S2**: Fluorescence microscopy time series of a Vero cell infected with RSV N(I183*), labeled with SiR-Tz (N; magenta) and Alexa488-D25 Fab (F; green), and imaged at 10-minute intervals over the course of 6h. An inclusion body that coalesces within the timelapse is indicated by the white circle. Images are displayed as a maximum intensity projection. Scale bar = 10 μm.

**Supplementary Movie S3**: Fluorescence microscopy time series tracking an inclusion body within a Vero cell infected with RSV N(I183*) and labeled with JF549-Tz. Images are collected at 10 s intervals over the course of 1h. Images show a single confocal slice displayed in inverted contrast. Scale bar = 1 μm.

**Supplementary Movie S4**: Fluorescence microscopy time series tracking an inclusion body within a Vero cell infected with RSV N(I183*) and labeled with JF549-Tz (N, shown in magenta) and Alexa488-D25 Fab (F, shown in green). Images are collected at 1-minute intervals over the course of 6h. Images show a single confocal slice. Scale bar = 1 μm.

**Sequences**

RSV A2 (Line 19F) genomic sequence:

NS1-IRES-mTagBFP2/NS2/N(I183*)/P/M/SH/G(ybbR)/(G5)F/M2/L

ACGCGAAAAAATGCGTACAACAAACTTGCATAAACCAAAAAAATGGGGCAAATAAGAATTTGATAAGTACCACTTAAATTTAACTCCCTTGGTTAGAGATGGGCAGCAATTC
ATTGAGTATGATAAAAGTTAGATTACAAAATTTGTTTGACAATGATGAAGTAGCATTGTTAAAAATAACATGCTATACTGATAAATTAATACATTTAACTAACGCTTTGGCT
AAGGCAGTGATACATACAATCAAATTGAATGGCATTGTGTTTGTGCATGTTATTACAAGTAGTGATATTTGCCCTAATAATAATATTGTAGTAAAATCCAATTTCACAACAA
TGCCAGTACTACAAAATGGAGGTTATATATGGAAATGATGGAATTAACACATTGCTCTCAACCTAATGGTCTACTAGATGACAATTGTGAAATTAAATTCTCCAAAAAACT
AAGTGATTCAACAATGACCAATTATATGAATCAATTATCTGAATTACTTGGATTTGATCTTAATCCATAACGGTTTCCCTCTAGCGGGATCAATTCCGCCCCCCCCCCCCTAA
CGTTACTGGCCGAAGCCGCTTGGAATAAGGCCGGTGTGCGTTTGTCTATATGTTATTTTCCACCATATTGCCGTCTTTTGGCAATGTGAGGGCCCGGAAACCTGGCCCTGTC
TTCTTGACGAGCATTCCTAGGGGTCTTTCCCCTCTCGCCAAAGGAATGCAAGGTCTGTTGAATGTCGTGAAGGAAGCAGTTCCTCTGGAAGCTTCTTGAAGACAAACAACGT
CTGTAGCGACCCTTTGCAGGCAGCGGAACCCCCCACCTGGCGACAGGTGCCTCTGCGGCCAAAAGCCACGTGTATAAGATACACCTGCAAAGGCGGCACAACCCCAGTGCCA
CGTTGTGAGTTGGATAGTTGTGGAAAGAGTCAAATGGCTCTCCTCAAGCGTATTCAACAAGGGGCTGAAGGATGCCCAGAAGGTACCCCATTGTATGGGATCTGATCTGGGG
CCTCGGTGCACATGCTTTACATGTGTTTAGTCGAGGTTAAAAAAACGTCTAGGCCCCCCGAACCACGGGGACGTGGTTTTCCTTTGAAAAACACGATAATACCATGAGCGAG
CTGATTAAGGAGAACATGCACATGAAGCTGTACATGGAGGGCACCGTGGACAACCATCACTTCAAGTGCACATCCGAGGGCGAAGGCAAGCCCTACGAGGGCACCCAGACCA
TGAGAATCAAGGTGGTCGAGGGCGGCCCTCTCCCCTTCGCCTTCGACATCCTGGCTACTAGCTTCCTCTACGGCAGCAAGACCTTCATCAACCACACCCAGGGCATCCCCGA
CTTCTTCAAGCAGTCCTTCCCTGAGGGCTTCACATGGGAGAGAGTCACCACATACGAAGACGGGGGCGTGCTGACCGCTACCCAGGACACCAGCCTCCAGGACGGCTGCCTC
ATCTACAACGTCAAGATCAGAGGGGTGAACTTCACATCCAACGGCCCTGTGATGCAGAAGAAAACACTCGGCTGGGAGGCCTTCACCGAGACGCTGTACCCCGCTGACGGCG
GCCTGGAAGGCAGAAACGACATGGCCCTGAAGCTCGTGGGCGGGAGCCATCTGATCGCAAACGCCAAGACCACATATAGATCCAAGAAACCCGCTAAGAACCTCAAGATGCC
TGGCGTCTACTATGTGGACTACAGACTGGAAAGAATCAAGGAGGCCAACAACGAGACCTACGTCGAGCAGCACGAGGTGGCAGTGGCCAGATACTGCGACCTCCCTAGCAAA
CTGGGGCACAAGCTTAATTAAATTATAATTAATATCAACTAGCAAATCAATGTCACTAACACCATTAGTTAATATAAAACTTAACAGAAGACAAAAATGGGGCAAATAAATC
AATTCAGCCAACCCAACCATGGACACAACCCACAATGATAATACACCACAAAGACTGATGATCACAGACATGAGACCGTTGTCACTTGAGACCATAATAACATCACTAACCA
GAGACATCATAACACACAAATTTATATACTTGATAAATCATGAATGCATAGTGAGAAAACTTGATGAAAAACAGGCCACATTTACATTCCTGGTCAACTATGAAATGAAACT
ATTACACAAAGTAGGAAGCACTAAATATAAAAAATATACTGAATACAACACAAAATATGGCACTTTCCCTATGCCAATATTCATCAATCATGATGGGTTCTTAGAATGCATT
GGCATTAAGCCTACAAAGCATACTCCCATAATATACAAGTATGATCTCAATCCATAAATTTCAACACAATATTCACACAATCTAAAACAACAACTCTATGCATAACTATACT
CCATAGTCCAGATGGAGCCTGAAAATTATAGTAATTTAAAATTAAGGAGAGATATAAGATAGAAGATGGGGCAAATACAAAGATGGCTCTTAGCAAAGTCAAGTTGAATGAT
ACACTCAACAAAGATCAACTTCTGTCATCCAGCAAATACACCATCCAACGGAGCACAGGAGATAGTATTGATACTCCTAATTATGATGTGCAGAAACACATCAATAAGTTAT
GTGGCATGTTATTAATCACAGAAGATGCTAATCATAAATTCACTGGGTTAATAGGTATGTTATATGCGATGTCTAGGTTAGGAAGAGAAGACACCATAAAAATACTCAGAGA
TGCGGGATATCATGTAAAAGCAAATGGAGTAGATGTAACAACACATCGTCAAGACATTAATGGAAAAGAAATGAAATTTGAAGTGTTAACATTGGCAAGCTTAACAACTGAA
ATTCAAATCAACATTGAGATAGAATCTAGAAAATCCTACAAAAAAATGCTAAAAGAAATGGGAGAGGTAGCTCCAGAATACAGGCATGACTCTCCTGATTGTGGGATGATAA
TATTATGTATAGCAGCATTAGTAATAACTAAATTAGCAGCAGGGGACAGATCTGGTCTTACCGCTGTATAGCGCAGGGCCAATAATGTCCTAAAAAATGAAATGAAACGTTA
CAAAGGCTTACTACCCAAGGACATAGCCAACAGCTTCTATGAAGTGTTTGAAAAACATCCCCACTTTATAGATGTTTTTGTTCATTTTGGTATAGCACAATCTTCTACCAGA
GGTGGCAGTAGAGTTGAAGGGATTTTTGCAGGATTGTTTATGAATGCCTATGGTGCAGGGCAAGTGATGTTACGGTGGGGAGTCTTAGCAAAATCAGTTAAAAATATTATGT
TAGGACATGCTAGTGTGCAAGCAGAAATGGAACAAGTTGTTGAGGTTTATGAATATGCCCAAAAATTGGGTGGTGAAGCAGGATTCTACCATATATTGAACAACCCAAAAGC
ATCATTATTATCTTTGACTCAATTTCCTCACTTCTCCAGTGTAGTATTAGGCAATGCTGCTGGCCTAGGCATAATGGGAGAGTACAGAGGTACACCGAGGAATCAAGATCTA
TATGATGCAGCAAAGGCATATGCTGAACAACTCAAAGAAAATGGTGTGATTAACTACAGTGTACTAGACTTGACAGCAGAAGAACTAGAGGCTATCAAACATCAGCTTAATC
CAAAAGATAATGATGTAGAGCTTTGAGTTAATAAAAAATGGGGCAAATAAATCATCATGGAAAAGTTTGCTCCTGAATTCCATGGAGAAGATGCAAACAACAGGGCTACTAA
ATTCCTAGAATCAATAAAGGGCAAATTCACATCACCCAAAGATCCCAAGAAAAAAGATAGTATCATATCTGTCAACTCAATAGATATAGAAGTAACCAAAGAAAGCCCTATA
ACATCAAATTCAACTATTATCAACCCAACAAATGAGACAGATGATACTGCAGGGAACAAGCCCAATTATCAAAGAAAACCTCTAGTAAGTTTCAAAGAAGACCCTACACCAA
GTGATAATCCCTTTTCTAAACTATACAAAGAAACCATAGAAACATTTGATAACAATGAAGAAGAATCCAGCTATTCATACGAAGAAATAAATGATCAGACAAACGATAATAT
AACAGCAAGATTAGATAGGATTGATGAAAAATTAAGTGAAATACTAGGAATGCTTCACACATTAGTAGTGGCAAGTGCAGGACCTACATCTGCTCGGGATGGTATAAGAGAT
GCCATGATTGGTTTAAGAGAAGAAATGATAGAAAAAATCAGAACTGAAGCATTAATGACCAATGACAGATTAGAAGCTATGGCAAGACTCAGGAATGAGGAAAGTGAAAAGA
TGGCAAAAGACACATCAGATGAAGTGTCTCTCAATCCAACATCAGAGAAATTGAACAACCTATTGGAAGGGAATGATAGTGACAATGATCTATCACTTGAAGATTTCTGATT
AGTTACCACTCTTCACATCAACACACAATACCAACAGAAGACCAACAAACTAACCAACCCAATCATCCAACCAAACATCCATCCGCCAATCAGCCAAACAGCCAACAAAACA
ACCAGCCAATCCAAAACTAACCACCCGGAAAAAATCTATAATATAGTTACAAAAAAAGGAAAGGGTGGGGCAAATATGGAAACATACGTGAACAAGCTTCACGAAGGCTCCA
CATACACAGCTGCTGTTCAATACAATGTCTTAGAAAAAGACGATGACCCTGCATCACTTACAATATGGGTGCCCATGTTCCAATCATCTATGCCAGCAGATTTACTTATAAA
AGAACTAGCTAATGTCAACATACTAGTGAAACAAATATCCACACCCAAGGGACCTTCACTAAGAGTCATGATAAACTCAAGAAGTGCAGTGCTAGCACAAATGCCCAGCAAA
TTTACCATATGCGCTAATGTGTCCTTGGATGAAAGAAGCAAACTAGCATATGATGTAACCACACCCTGTGAAATCAAGGCATGTAGTCTAACATGCCTAAAATCAAAAAATA
TGTTGACTACAGTTAAAGATCTCACTATGAAGACACTCAACCCTACACATGATATTATTGCTTTATGTGAATTTGAAAACATAGTAACATCAAAAAAAGTCATAATACCAAC
ATACCTAAGATCCATCAGTGTCAGAAATAAAGATCTGAACACACTTGAAAATATAACAACCACTGAATTCAAAAATGCTATCACAAATGCAAAAATCATCCCTTACTCAGGA
TTACTATTAGTCATCACAGTGACTGACAACAAAGGAGCATTCAAATACATAAAGCCACAAAGTCAATTCATAGTAGATCTTGGAGCTTACCTAGAAAAAGAAAGTATATATT
ATGTTACCACAAATTGGAAGCACACAGCTACACGATTTGCAATCAAACCCATGGAAGATTAACCTTTTTCCTCTACATCAGTGTGTTAATTCATACAAACTTTCTACCTACA
TTCTTCACTTCACCATCACAATCACAAACACTCTGTGGTTCAACCAATCAAACAAAACTTATCTGAAGTCCCAGATCATCCCAAGTCATTGTTTATCAGATCTAGTACTCAA
ATAAGTTAATAAAAAATATACACATGGGGCAAATAATCATTGGAGGAAATCCAACTAATCACAATATCTGTTAACATAGACAAGTCCACACACCATACAGAATCAACCAATG
GAAAATACATCCATAACAATAGAATTCTCAAGCAAATTCTGGCCTTACTTTACACTAATACACATGATCACAACAATAATCTCTTTGCTAATCATAATCTCCATCATGATTG
CAATACTAAACAAACTTTGTGAATATAACGTATTCCATAACAAAACCTTTGAGTTACCAAGAGCTCGAGTCAACACATAGCATTCATCAATCCAACAGCCCAAAACAGTAAC
CTTGCATTTAAAAATGAACAACCCCTACCTCTTTACAACACCTCATTAACATCCCACCATGCAAACCACTATCCATACTATAAAGTAGTTAATTAAAAATAGTCATAACAAT
GAACTAGGATATCAAGACTAACAATAACATTGGGGCAAATGCAAACATGTCCAAAAACAAGGACCAACGCACCGCTAAGACATTAGAAAGGACCTGGGACACTCTCAATCAT
TTATTATTCATATCATCGTGCTTATATAAGTTAAATCTTAAATCTGTAGCACAAATCACATTATCCATTCTGGCAATGATAATCTCAACTTCACTTATAATTGCAGCCATCA
TATTCATAGCCTCGGCAAACCACAAAGTCACACCAACAACTGCAATCATACAAGATGCAACAAGCCAGATCAAGAACACAACCCCAACATACCTCACCCAGAATCCTCAGCT
TGGAATCAGTCCCTCTAATCCGTCTGAAATTACATCACAAATCACCACCATACTAGCTTCAACAACACCAGGAGTCAAGTCAACCCTGCAATCCACAACAGTCAAGACCAAA
AACACAACAACAACTCAAACACAACCCAGCAAGCCCACCACAAAACAACGCCAAAACAAACCACCAAGCAAACCCAATAATGATTTTCACTTTGAAGTGTTCAACTTTGTAC
CCTGCAGCATATGCAGCAACAATCCAACCTGCTGGGCTATCTGCAAAAGAATACCAAACAAAAAACCAGGAAAGAAAACCACTACCAAGCCCACAAAAAAACCAACCCTCAA
GACAACCAAAAAAGATCCCAAACCTCAAACCACTAAATCAAAGGAAGTACCCACCACCAAGCCCACAGAAGAGCCAACCATCAACACCACCAAAACAAACATCATAACTACA
CTACTCACCTCCAACACCACAGGAAATCCAGAACTCACAAGTCAAATGGAAACCTTCCACTCAACTTCCTCCGAAGGCAATCCAAGCCCTTCTCAAGTCTCTACAACATCCG
AGTACCCATCACAACCTTCATCTCCACCCAACACACCACGCCAG**GGCGGCGGCGATTCTCTTGAATTTATTGCTAGTAAGCTTGCG**TAGTTACTTAAAAACATATTATCACA
AAAAGCCATGACCAACTTAAACAGAATCAAAATAAACTCTGGGGCAAATAACAATGGAGTTGCCAATCCTCAAAGCAAATGCAATTACCACAATCCTCGCTGCAGTCACATT
TTGCTTTGCTTCTGGT**GGCGGAGGAGGGGGC**CAAAACATCACTGAAGAATTTTATCAATCAACATGCAGTGCAGTTAGCAAAGGCTATCTTAGTGCTCTAAGAACTGGTTGG
TATACTAGTGTTATAACTATAGAATTAAGTAATATCAAGAAAAATAAGTGTAATGGAACAGATGCTAAGGTAAAATTGATGAAACAAGAATTAGATAAATATAAAAATGCTG
TAACAGAATTGCAGTTGCTCATGCAAAGCACACCAGCAGCAAACAATCGAGCCAGAAGAGAACTACCAAGGTTTATGAATTATACACTCAACAATACCAAAAAAACCAATGT
AACATTAAGCAAGAAAAGGAAAAGAAGATTTCTTGGTTTTTTGTTAGGTGTTGGATCTGCAATCGCCAGTGGCATTGCTGTATCTAAGGTCCTGCACTTAGAAGGAGAAGTG
AACAAGATCAAAAGTGCTCTACTATCCACAAACAAGGCCGTAGTCAGCTTATCAAATGGAGTTAGTGTCTTAACCAGCAGAGTGTTAGACCTCAAAAACTATATAGATAAAC
AATTGTTACCTATTGTGAATAAGCAAAGCTGCAGAATATCAAATATAGAAACTGTGATAGAGTTCCAACAAAAGAACAACAGACTACTAGAGATTACCAGGGAATTTAGTGT

TAATGCAGGTGTAACTACACCTGTAAGCACTTACATGTTAACTAATAGTGAATTATTGTCATTAATCAATGATATGCCTATAACAAATGATCAGAAAAAGTTAATGTCCAAC
AATGTTCAAATAGTTAGACAGCAAAGTTACTCTATCATGTCCATAATAAAAGAGGAAGTCTTAGCATATGTAGTACAATTACCACTATATGGTGTGATAGATACACCTTGTT
GGAAATTACACACATCCCCTCTATGTACAACCAACACAAAAGAAGGGTCAAACATCTGTTTAACAAGAACTGACAGAGGATGGTACTGTGACAATGCAGGATCAGTATCTTT
CTTCCCACAAGCTGAAAAATGTAAAGTTCAATCGAATCGAGTATTTTGTGACACAATGTACAGTTTAACATTACCAAGTGAAGTAAATCTCTGCAATGTTGACATATTCAAT
CCCAAATATGATTGTAAAATTATGACTTCAAAAACAGATGTAAGCAGCTCCGTTATCACATCTCTAGGAGCCATTGTGTCATGCTATGGCAAAACTAAATGTACAGCATCCA
ATAAAAATCGTGGAATCATAAAGACATTTTCTAACGGGTGTGATTATGTATCAAATAAAGGGGTGGACACTGTGTCTGTAGGTAACACATTATATTATGTAAATAAGCAAGA
AGGCAAAAGTCTCTATGTAAAAGGTGAACCAATAATAAATTTCTATGACCCATTAGTATTCCCCTCTGATGAATTTGATGCATCAATATCTCAAGTCAATGAGAAGATTAAC
CAGAGTTTAGCATTTATTCGTAAATCCGATGAATTATTACATAATGTAAATGCTGGTAAATCAACCACAAATATCATGATAACTACTATAATTATAGTGATTATAGTAATAT
TGTTATCATTAATTGCTGTTGGACTGCTCCTATACTGTAAGGCCAGAAGCACACCAATCACACTAAGCAAGGATCAACTGAGTGGTATAAATAATATTGCATTTAGTAACTG
AATAAAAATAGCACCTAATCATGTTCTTACAATGGTTTACTATCTGCTCATAGACAACCCATCTATCATTGGATTTTCTTAAAATCTGAACTTCATCGAAACTCTTATCTAT
AAACCATCTCACTTACACTATTTAAGTAGATTCCTAGTTTATAGTTATATAAAACACAATTGAATGCCAGATTAACTTACCATCTGTAAAAATGAAAACTGGGGCAAATATG
TCACGAAGGAATCCTTGCAAATTTGAAATTCGAGGTCATTGCTTAAATGGTAAGAGGTGTCATTTTAGTCATAATTATTTTGAATGGCCACCCCATGCACTGCTTGTAAGAC
AAAACTTTATGTTAAACAGAATACTTAAGTCTATGGATAAAAGTATAGATACCTTATCAGAAATAAGTGGAGCTGCAGAGTTGGACAGAACAGAAGAGTATGCTCTTGGTGT
AGTTGGAGTGCTAGAGAGTTATATAGGATCAATAAACAATATAACTAAACAATCAGCATGTGTTGCCATGAGCAAACTCCTCACTGAACTCAATAGTGATGATATCAAAAAG
CTGAGGGACAATGAAGAGCTAAATTCACCCAAGATAAGAGTGTACAATACTGTCATATCATATATTGAAAGCAACAGGAAAAACAATAAACAAACTATCCATCTGTTAAAAA
GATTGCCAGCAGACGTATTGAAGAAAACCATCAAAAACACATTGGATATCCATAAGAGCATAACCATCAACAACCCAAAAGAATCAACTGTTAGTGATACAAATGACCATGC
CAAAAATAATGATACTACCTGACAAATATCCTTGTAGTATAACTTCCATACTAATAACAAGTAGATGTAGAGTTACTATGTATAATCAAAAGAACACACTATATTTCAATCA
AAACAACCCAAATAACCATATGTACTCACCGAATCAAACATTCAATGAAATCCATTGGACCTCTCAAGAATTGATTGACACAATTCAAATTTTTCTACAACATCTAGGTATT
ATTGAGGATATATATACAATATATATATTAGTGTCATAACACTCAATTCTAACACTCACCACATCGTTACATTATTAATTCAAACAATTCAAGTTGTGGGACAAAATGGATC
CCATTATTAATGGAAATTCTGCTAATGTTTATCTAACCGATAGTTATTTAAAAGGTGTTATCTCTTTCTCAGAGTGTAATGCTTTAGGAAGTTACATATTCAATGGTCCTTA
TCTCAAAAATGATTATACCAACTTAATTAGTAGACAAAATCCATTAATAGAACACATGAATCTAAAGAAACTAAATATAACACAGTCCTTAATATCTAAGTATCATAAAGGT
GAAATAAAATTAGAAGAACCTACTTATTTTCAGTCATTACTTATGACATACAAGAGTATGACCTCGTCAGAACAGATTGCTACCACTAATTTACTTAAAAAGATAATAAGAA
GAGCTATAGAAATAAGTGATGTCAAAGTCTATGCTATATTGAATAAACTAGGGCTTAAAGAAAAGGACAAGATTAAATCCAACAATGGACAAGATGAAGACAACTCAGTTAT
TACGACCATAATCAAAGATGATATACTTTCAGCTGTTAAAGATAATCAATCTCATCTTAAAGCAGACAAAAATCACTCTACAAAACAAAAAGACACAATCAAAACAACACTC
TTGAAGAAATTGATGTGTTCAATGCAACATCCTCCATCATGGTTAATACATTGGTTTAACTTATACACAAAATTAAACAACATATTAACACAGTATCGATCAAATGAGGTAA
AAAACCATGGGTTTACATTGATAGATAATCAAACTCTTAGTGGATTTCAATTTATTTTGAACCAATATGGTTGTATAGTTTATCATAAGGAACTCAAAAGAATTACTGTGAC
AACCTATAATCAATTCTTGACATGGAAAGATATTAGCCTTAGTAGATTAAATGTTTGTTTAATTACATGGATTAGTAACTGCTTGAACACATTAAATAAAAGCTTAGGCTTA
AGATGCGGATTCAATAATGTTATCTTGACACAACTATTCCTTTATGGAGATTGTATACTAAAGCTATTTCACAATGAGGGGTTCTACATAATAAAAGAGGTAGAGGGATTTA
TTATGTCTCTAATTTTAAATATAACAGAAGAAGATCAATTCAGAAAACGATTTTATAATAGTATGCTCAACAACATCACAGATGCTGCTAATAAAGCTCAGAAAAATCTGCT
ATCAAGAGTATGTCATACATTATTAGATAAGACAGTGTCCGATAATATAATAAATGGCAGATGGATAATTCTATTAAGTAAGTTCCTTAAATTAATTAAGCTTGCAGGTGAC
AATAACCTTAACAATCTGAGTGAACTATATTTTTTGTTCAGAATATTTGGACACCCAATGGTAGATGAAAGACAAGCCATGGATGCTGTTAAAATTAATTGCAATGAGACCA
AATTTTACTTGTTAAGCAGTCTGAGTATGTTAAGAGGTGCCTTTATATATAGAATTATAAAAGGGTTTGTAAATAATTACAACAGATGGCCTACTTTAAGAAATGCTATTGT
TTTACCCTTAAGATGGTTAACTTACTATAAACTAAACACTTATCCTTCTTTGTTGGAACTTACAGAAAGAGATTTGATTGTGTTATCAGGACTACGTTTCTATCGTGAGTTT
CGGTTGCCTAAAAAAGTGGATCTTGAAATGATTATAAATGATAAAGCTATATCACCTCCTAAAAATTTGATATGGACTAGTTTCCCTAGAAATTACATGCCATCACACATAC
AAAACTATATAGAACATGAAAAATTAAAATTTTCCGAGAGTGATAAATCAAGAAGAGTATTAGAGTATTATTTAAGAGATAACAAATTCAATGAATGTGATTTATACAACTG
TGTAGTTAATCAAAGTTATCTCAACAACCCTAATCATGTGGTATCATTGACAGGCAAAGAAAGAGAACTCAGTGTAGGTAGAATGTTTGCAATGCAACCGGGAATGTTCAGA
CAGGTTCAAATATTGGCAGAGAAAATGATAGCTGAAAACATTTTACAATTCTTTCCTGAAAGTCTTACAAGATATGGTGATCTAGAACTACAAAAAATATTAGAATTGAAAG
CAGGAATAAGTAACAAATCAAATCGCTACAATGATAATTACAACAATTACATTAGTAAGTGCTCTATCATCACAGATCTCAGCAAATTCAATCAAGCATTTCGATATGAAAC
GTCATGTATTTGTAGTGATGTGCTGGATGAACTGCATGGTGTACAATCTCTATTTTCCTGGTTACATTTAACTATTCCTCATGTCACAATAATATGCACATATAGGCATGCA
CCCCCCTATATAGGAGATCATATTGTAGATCTTAACAATGTAGATGAACAAAGTGGATTATATAGATATCACATGGGTGGCATCGAAGGGTGGTGTCAAAAACTGTGGACCA
TAGAAGCTATATCACTATTGGATCTAATATCTCTCAAAGGGAAATTCTCAATTACTGCTTTAATTAATGGTGACAATCAATCAATAGATATAAGCAAACCAATCAGACTCAT
GGAAGGTCAAACTCATGCTCAAGCAGATTATTTGCTAGCATTAAATAGCCTTAAATTACTGTATAAAGAGTATGCAGGCATAGGCCACAAATTAAAAGGAACTGAGACTTAT
ATATCACGAGATATGCAATTTATGAGTAAAACAATTCAACATAACGGTGTATATTACCCAGCTAGTATAAAGAAAGTCCTAAGAGTGGGACCGTGGATAAACACTATACTTG
ATGATTTCAAAGTGAGTCTAGAATCTATAGGTAGTTTGACACAAGAATTAGAATATAGAGGTGAAAGTCTATTATGCAGTTTAATATTTAGAAATGTATGGTTATATAATCA
GATTGCTCTACAATTAAAAAATCATGCATTATGTAACAATAAACTATATTTGGACATATTAAAGGTTCTGAAACACTTAAAAACCTTTTTTAATCTTGATAATATTGATACA
GCATTAACATTGTATATGAATTTACCCATGTTATTTGGTGGTGGTGATCCCAACTTGTTATATCGAAGTTTCTATAGAAGAACTCCTGACTTCCTCACAGAGGCTATAGTTC
ACTCTGTGTTCATACTTAGTTATTATACAAACCATGACTTAAAAGATAAACTTCAAGATCTGTCAGATGATAGATTGAATAAGTTCTTAACATGCATAATCACGTTTGACAA
AAACCCTAATGCTGAATTCGTAACATTGATGAGAGATCCTCAAGCTTTAGGGTCTGAGAGACAAGCTAAAATTACTAGCGAAATCAATAGACTGGCAGTTACAGAGGTTTTG
AGTACAGCTCCAAACAAAATATTCTCCAAAAGTGCACAACATTATACTACTACAGAGATAGATCTAAATGATATTATGCAAAATATAGAACCTACATATCCTCATGGGCTAA
GAGTTGTTTATGAAAGTTTACCCTTTTATAAAGCAGAGAAAATAGTAAATCTTATATCAGGTACAAAATCTATAACTAACATACTGGAAAAAACTTCTGCCATAGACTTAAC
AGATATTGATAGAGCCACTGAGATGATGAGGAAAAACATAACTTTGCTTATAAGGATACTTCCATTGGATTGTAACAGAGATAAAAGAGAGATATTGAGTATGGAAAACCTA
AGTATTACTGAATTAAGCAAATATGTTAGGGAAAGATCTTGGTCTTTATCCAATATAGTTGGTGTTACATCACCCAGTATCATGTATACAATGGACATCAAATATACTACAA
GCACTATATCTAGTGGCATAATTATAGAGAAATATAATGTTAACAGTTTAACACGTGGTGAGAGAGGACCCACTAAACCATGGGTTGGTTCATCTACACAAGAGAAAAAAAC
AATGCCAGTTTATAATAGACAAGTCTTAACCAAAAAACAGAGAGATCAAATAGATCTATTAGCAAAATTGGATTGGGTGTATGCATCTATAGATAACAAGGATGAATTCATG
GAAGAACTCAGCATAGGAACCCTTGGGTTAACATATGAAAAGGCCAAGAAATTATTTCCACAATATTTAAGTGTCAATTATTTGCATCGCCTTACAGTCAGTAGTAGACCAT
GTGAATTCCCTGCATCAATACCAGCTTATAGAACAACAAATTATCACTTTGACACTAGCCCTATTAATCGCATATTAACAGAAAAGTATGGTGATGAAGATATTGACATAGT
ATTCCAAAACTGTATAAGCTTTGGCCTTAGTTTAATGTCAGTAGTAGAACAATTTACTAATGTATGTCCTAACAGAATTATTCTCATACCTAAGCTTAATGAGATACATTTG
ATGAAACCTCCCATATTCACAGGTGATGTTGATATTCACAAGTTAAAACAAGTGATACAAAAACAGCATATGTTTTTACCAGACAAAATAAGTTTGACTCAATATGTGGAAT
TATTCTTAAGTAATAAAACACTCAAATCTGGATCTCATGTTAATTCTAATTTAATATTGGCACATAAAATATCTGACTATTTTCATAATACTTACATTTTAAGTACTAATTT
AGCTGGACATTGGATTCTGATTATACAACTTATGAAAGATTCTAAAGGTATTTTTGAAAAAGATTGGGGAGAGGGATATATAACTGATCATATGTTTATTAATTTGAAAGTT
TTCTTCAATGCTTATAAGACCTATCTCTTGTGTTTTCATAAAGGTTATGGCAAAGCAAAGCTGGAGTGTGATATGAACACTTCAGATCTTCTATGTGTATTGGAATTAATAG
ACAGTAGTTATTGGAAGTCTATGTCTAAGGTATTTTTAGAACAAAAAGTTATCAAATACATTCTTAGCCAAGATGCAAGTTTACATAGAGTAAAAGGATGTCATAGCTTCAA
ATTATGGTTTCTTAAACGTCTTAATGTAGCAGAATTCACAGTTTGCCCTTGGGTTGTTAACATAGATTATCATCCAACACATATGAAAGCAATATTAACTTATATAGATCTT
GTTAGAATGGGATTGATAAATATAGATAGAATACACATTAAAAATAAACACAAATTCAATGATGAATTTTATACTTCTAATCTCTTCTACATTAATTATAACTTCTCAGATA
ATACTCATCTATTAACTAAACATATAAGGATTGCTAATTCTGAATTAGAAAATAATTACAACAAATTATATCATCCTACACCAGAAACCCTAGAGAATATACTAGCCAATCC
GATTAAAAGTAATGACAAAAAGACACTGAATGACTATTGTATAGGTAAAAATGTTGACTCAATAATGTTACCATTGTTATCTAATAAGAAGCTTATTAAATCGTCTGCAATG
ATTAGAACCAATTACAGCAAACAAGATTTGTATAATTTATTCCCTATGGTTGTGATTGATAGAATTATAGATCATTCAGGCAATACAGCCAAATCCAACCAACTTTACACTA
CTACTTCCCACCAAATATCTTTAGTGCACAATAGCACATCACTTTACTGCATGCTTCCTTGGCATCATATTAATAGATTCAATTTTGTATTTAGTTCTACAGGTTGTAAAAT
TAGTATAGAGTATATTTTAAAAGATCTTAAAATTAAAGATCCCAATTGTATAGCATTCATAGGTGAAGGAGCAGGGAATTTATTATTGCGTACAGTAGTGGAACTTCATCCT
GACATAAGATATATTTACAGAAGTCTGAAAGATTGCAATGATCATAGTTTACCTATTGAGTTTTTAAGGCTGTACAATGGACATATCAACATTGATTATGGTGAAAATTTGA
CCATTCCTGCTACAGATGCAACCAACAACATTCATTGGTCTTATTTACATATAAAGTTTGCTGAACCTATCAGTCTTTTTGTCTGTGATGCCGAATTGTCTGTAACAGTCAA
CTGGAGTAAAATTATAATAGAATGGAGCAAGCATGTAAGAAAGTGCAAGTACTGTTCCTCAGTTAATAAATGTATGTTAATAGTAAAATATCATGCTCAAGATGATATTGAT
TTCAAATTAGACAATATAACTATATTAAAAACTTATGTATGCTTAGGCAGTAAGTTAAAGGGATCGGAGGTTTACTTAGTCCTTACAATAGGTCCTGCGAATATATTCCCAG
TATTTAATGTAGTACAAAATGCTAAATTGATACTATCAAGAACCAAAAATTTCATCATGCCTAAGAAAGCTGATAAAGAGTCTATTGATGCAAATATTAAAAGTTTGATACC
CTTTCTTTGTTACCCTATAACAAAAAAAGGAATTAATACTGCATTGTCAAAACTAAAGAGTGTTGTTAGTGGAGATATACTATCATATTCTATAGCTGGACGTAATGAAGTT
TTCAGCAATAAACTTATAAATCATAAGCATATGAACATCTTAAAATGGTTCAATCATGTTTTAAATTTCAGATCAACAGAACTAAACTATAACCATTTATATATGGTAGAAT

CTACATATCCTTACCTAAGTGAATTGTTAAACAGCTTGACAACCAATGAACTTAAAAAACTGATTAAAATCACAGGTAGTCTGTTATACAACTTTCATAATGAATAATGAAT
AAAGATCTTATAATAAAAATTCCCATAGCTATACACTAACACTGTATTCAATTATAGTTATTAAAAATTAAAAATCATATAATTTTTTAAATAACTTTTAGTGAACTAATCC
TAAAGTTATCATTTTAATCTTGGAGGAATAAATTTAAACCCTAATCTAATTGGTTTATATGTGTATTAACTAAATTACGAGATATTAGTTTTTGACACTTTTTTTCTCGT

BFP mini-genome (inserted into pcDNA3.1(+) between BglII and BbsI sites)

T7 promoter/Hammerhead ribozyme/Leader/mTagBFP2/Trailer/HDV ribozyme/T7 terminator

GACGGATCGGGAGATCTTTCGAAACGAAATTAATACGACTCACTATAGGGAAGTTTTTCGCGTCTGATGAGGCCGTTAGGCCGAAACTCCTCTCCGGAGTCACGCGAAAAAA
TGCGTACAACAAACTTGCATAAACCAAAAAAATGGGGCAAATAAGAATTTGATAAGTACCACTTAAATTTAACTCCCTTGGTTAGAGATGAGCGAGCTGATTAAGGAGAACA
TGCACATGAAGCTGTACATGGAGGGCACCGTGGACAACCATCACTTCAAGTGCACATCCGAGGGCGAAGGCAAGCCCTACGAGGGCACCCAGACCATGAGAATCAAGGTGGT
CGAGGGCGGCCCTCTCCCCTTCGCCTTCGACATCCTGGCTACTAGCTTCCTCTACGGCAGCAAGACCTTCATCAACCACACCCAGGGCATCCCCGACTTCTTCAAGCAGTCC
TTCCCTGAGGGCTTCACATGGGAGAGAGTCACCACATACGAAGACGGGGGCGTGCTGACCGCTACCCAGGACACCAGCCTCCAGGACGGCTGCCTCATCTACAACGTCAAGA
TCAGAGGGGTGAACTTCACATCCAACGGCCCTGTGATGCAGAAGAAAACACTCGGCTGGGAGGCCTTCACCGAGACGCTGTACCCCGCTGACGGCGGCCTGGAAGGCAGAAA
CGACATGGCCCTGAAGCTCGTGGGCGGGAGCCATCTGATCGCAAACGCCAAGACCACATATAGATCCAAGAAACCCGCTAAGAACCTCAAGATGCCTGGCGTCTACTATGTG
GACTACAGACTGGAAAGAATCAAGGAGGCCAACAACGAGACCTACGTCGAGCAGCACGAGGTGGCAGTGGCCAGATACTGCGACCTCCCTAGCAAACTGGGGCACAAGCTTA
ATTAAATTATAATTAATATCAACTAGCAAATCAATGTCACTAACACCATTAGTTAATATAAAACTTAACAGAAGACAAAAATTTAAAAATCATATAATTTTTTAAATAACTT
TTAGTGAACTAATCCTAAAGTTATCATTTTAATCTTGGAGGAATAAATTTAAACCCTAATCTAATTGGTTTATATGTGTATTAACTAAATTACGAGATATTAGTTTTTGACA
CTTTTTTTCTCGTGGGTCGGCATGGCATCTCCACCTCCTCGCGGTCCGACCTGGGCATCCGAAGGAGGACGCACGTCCACTCGGATGGCTAAGGGAGAGCCACTGCTAACAA
AGCCCGAAAGGAAGCTGAGTTGGCTGCTGCCACCGCTGAGCAATAACTAGCATAACCCCTTGGGGCCTCTAAACGGGTCTTGAGGGGTTTTTTGCTGAAAGGAGGAACTATA
TACGCGTAGGCATGCTGGGGATGCGGTGGGCTCTATGGCTTCTGAGGCGGAAAGAACCAGCTGGGGCTCTAGGGGGTATCCCCACGCGCCCTGTAGCGGCGCATTAAGCGCG
GCGGGTGTGGTGGTTACGCGCAGCGTGACCGCTACACTTGCCAGCGCCCTAGCGCCCGCTCCTTTCGCTTTCTTCCCTTCCTTTCTCGCCACGTTCGCCGGCTTTCCCCGTC
AAGCTCTAAATCGGGGGCTCCCTTTAGGGTTCCGATTTAGTGCTTTACGGCACCTCGACCCCAAAAAACTTGATTAGGGTGATGGTTCACGTAGTGGGCCATCGCCCTGATA
GACGGTTTTTCGCCCTTTGACGTTGGAGTCCACGTTCTTTAATAGTGGACTCTTGTTCCAAACTGGAACAACACTCAACCCTATCTCGGTCTATTCTTTTGATTTATAAGGG
ATTTTGCCGATTTCGGCCTATTGGTTAAAAAATGAGCTGATTTAACAAAAATTTAACGCGAATTAATTCTGTGGAATGTGTGTCAGTTAGGGTGTGGAAAGTCCCCAGGCTC
CCCAGCAGGCAGAAGTATGCAAAGCATGCATCTCAATTAGTCAGCAACCAGGTGTGGAAAGTCCCCAGGCTCCCCAGCAGGCAGAAGTATGCAAAGCATGCATCTCAATTAG
TCAGCAACCATAGTCCCGCCCCTAACTCCGCCCATCCCGCCCCTAACTCCGCCCAGTTCCGCCCATTCTCCGCCCCATGGCTGACTAATTTTTTTTATTTATGCAGAGGCCG
AGGCCGCCTCTGCCTCTGAGCTATTCCAGAAGTAGTGAGGAGGCTTTTTTGGAGGCCTAGGCTTTTGCAAAAAGCTCCCGGGAGCTTGTATATCCATTTTCGGATCTGATCA
AGAGACAGGATGAGGATCGTTTCGCATGATTGAACAAGATGGATTGCACGCAGGTTCTCCGGCCGCTTGGGTGGAGAGGCTATTCGGCTATGACTGGGCACAACAGACAATC
GGCTGCTCTGATGCCGCCGTGTTCCGGCTGTCAGCGCAGGGGCGCCCGGTTCTTTTTGTCAAGACCGACCTGTCCGGTGCCCTGAATGAACTGCAGGACGAGGCAGCGCGGC
TATCGTGGCTGGCCACGACGGGCGTTCCTTGCGCAGCTGTGCTCGACGTTGTCACTGAAGCGGGAAGGGACTGGCTGCTATTGGGCGAAGTGCCGGGGCAGGATCTCCTGTC
ATCTCACCTTGCTCCTGCCGAGAAAGTATCCATCATGGCTGATGCAATGCGGCGGCTGCATACGCTTGATCCGGCTACCTGCCCATTCGACCACCAAGCGAAACATCGCATC
GAGCGAGCACGTACTCGGATGGAAGCCGGTCTTGTCGATCAGGATGATCTGGACGAAGAGCATCAGGGGCTCGCGCCAGCCGAACTGTTCGCCAGGCTCAAGGCGCGCATGC
CCGACGGCGAGGATCTCGTCGTGACCCATGGCGATGCCTGCTTGCCGAATATCATGGTGGAAAATGGCCGCTTTTCTGGATTCATCGACTGTGGCCGGCTGGGTGTGGCGGA
CCGCTATCAGGACATAGCGTTGGCTACCCGTGATATTGCTGAAGAGCTTGGCGGCGAATGGGCTGACCGCTTCCTCGTGCTTTACGGTATCGCCGCTCCCGATTCGCAGCGC
ATCGCCTTCTATCGCCTTCTTGACGAGTTCTTCTGAGCGGGACTCTGGGGTTCGAAATGACCGACCAAGCGACGCCCAACCTGCCATCACGAGATTTCGATTCCACCGCCGC
CTTCTATGAAAGGTTGGGCTTCGGAATCGTTTTCCGGGACGCCGGCTGGATGATCCTCCAGCGCGGGGATCTCATGCTGGAGTTCTTCGCCCACCCCAACTTGTTTATTGCA
GCTTATAATGGTTACAAATAAAGCAATAGCATCACAAATTTCACAAATAAAGCATTTTTTTCACTGCATTCTAGTTGTGGTTTGTCCAAACTCATCAATGTATCTTATCATG
TCTGTATACCGTCGACCTCTAGCTAGAGCTTGGCGTAATCATGGTCATAGCTGTTTCCTGTGTGAAATTGTTATCCGCTCACAATTCCACACAACATACGAGCCGGAAGCAT
AAAGTGTAAAGCCTGGGGTGCCTAATGAGTGAGCTAACTCACATTAATTGCGTTGCGCTCACTGCCCGCTTTCCAGTCGGGAAACCTGTCGTGCCAGCTGCATTAATGAATC
GGCCAACGCGCGGGGAGAGGCGGTTTGCGTATTGGGCGCTCTTCCGCTTCCTCGCTCACTGACTCGCTGCGCTCGGTCGTTCGGCTGCGGCGAGCGGTATCAGCTCACTCAA
AGGCGGTAATACGGTTATCCACAGAATCAGGGGATAACGCAGGAAAGAACATGTGAGCAAAAGGCCAGCAAAAGGCCAGGAACCGTAAAAAGGCCGCGTTGCTGGCGTTTTT
CCATAGGCTCCGCCCCCCTGACGAGCATCACAAAAATCGACGCTCAAGTCAGAGGTGGCGAAACCCGACAGGACTATAAAGATACCAGGCGTTTCCCCCTGGAAGCTCCCTC
GTGCGCTCTCCTGTTCCGACCCTGCCGCTTACCGGATACCTGTCCGCCTTTCTCCCTTCGGGAAGCGTGGCGCTTTCTCATAGCTCACGCTGTAGGTATCTCAGTTCGGTGT
AGGTCGTTCGCTCCAAGCTGGGCTGTGTGCACGAACCCCCCGTTCAGCCCGACCGCTGCGCCTTATCCGGTAACTATCGTCTTGAGTCCAACCCGGTAAGACACGACTTATC
GCCACTGGCAGCAGCCACTGGTAACAGGATTAGCAGAGCGAGGTATGTAGGCGGTGCTACAGAGTTCTTGAAGTGGTGGCCTAACTACGGCTACACTAGAAGAACAGTATTT
GGTATCTGCGCTCTGCTGAAGCCAGTTACCTTCGGAAAAAGAGTTGGTAGCTCTTGATCCGGCAAACAAACCACCGCTGGTAGCGGTTTTTTTGTTTGCAAGCAGCAGATTA
CGCGCAGAAAAAAAGGATCTCAAGAAGATCCTTTGATCTTTTCTACGGGGTCTGACGCTCAGTGGAACGAAAACTCACGTTAAGGGATTTTGGTCATGAGATTATCAAAAAG
GATCTTCACCTAGATCCTTTTAAATTAAAAATGAAGTTTTAAATCAATCTAAAGTATATATGAGTAAACTTGGTCTGACAGTTACCAATGCTTAATCAGTGAGGCACCTATC
TCAGCGATCTGTCTATTTCGTTCATCCATAGTTGCCTGACTCCCCGTCGTGTAGATAACTACGATACGGGAGGGCTTACCATCTGGCCCCAGTGCTGCAATGATACCGCGAG
ACCCACGCTCACCGGCTCCAGATTTATCAGCAATAAACCAGCCAGCCGGAAGGGCCGAGCGCAGAAGTGGTCCTGCAACTTTATCCGCCTCCATCCAGTCTATTAATTGTTG
CCGGGAAGCTAGAGTAAGTAGTTCGCCAGTTAATAGTTTGCGCAACGTTGTTGCCATTGCTACAGGCATCGTGGTGTCACGCTCGTCGTTTGGTATGGCTTCATTCAGCTCC
GGTTCCCAACGATCAAGGCGAGTTACATGATCCCCCATGTTGTGCAAAAAAGCGGTTAGCTCCTTCGGTCCTCCGATCGTTGTCAGAAGTAAGTTGGCCGCAGTGTTATCAC
TCATGGTTATGGCAGCACTGCATAATTCTCTTACTGTCATGCCATCCGTAAGATGCTTTTCTGTGACTGGTGAGTACTCAACCAAGTCATTCTGAGAATAGTGTATGCGGCG
ACCGAGTTGCTCTTGCCCGGCGTCAATACGGGATAATACCGCGCCACATAGCAGAACTTTAAAAGTGCTCATCATTGGAAAACGTTCTTCGGGGCGAAAACTCTCAAGGATC
TTACCGCTGTTGAGATCCAGTTCGATGTAACCCACTCGTGCACCCAACTGATCTTCAGCATCTTTTACTTTCACCAGCGTTTCTGGGTGAGCAAAAACAGGAAGGCAAAATG
CCGCAAAAAAGGGAATAAGGGCGACACGGAAATGTTGAATACTCATACTCTTCCTTTTTCAATATTATTGAAGCATTTATCAGGGTTATTGTCTCATGAGCGGATACATATT
TGAATGTATTTAGAAAAATAAACAAATAGGGGTTCCGCGCACATTTCCCCGAAAAGTGCCACCTGACGTC

piggyBac M15 PylT/RS vector sequence:

piggyBac inverted repeats hEF1a-HTLV promoter Flag-PylRS U6 tRNA-M15

GAACACGCAGCTAGATTAACCCTAGAAAGATAATCATATTGTGACGTACGTTAAAGATAATCATGTGTAAAATTGACGCATGTGTTTTATCGGTCTGTATATCGAGGTTTAT
TTATTAATTTGAATAGATATTAAGTTTTATTATATTTACACTTACATACTAATAATAAATTCAACAAACAATTTATTTATGTTTATTTATTTATTAAAAAAAAACAAAAACTC
AAAATTTCTTCTATAAAGTAACAAAACTTTTATGAGGGACAGCCCCCCCCCAAAGCCCCCAGGGATGTAATTACGTCCCTCCCCCGCTAGGGGGCAGCAGCGAGCCGCCCGG
GGCTCCGCTCCGGTCCGGCGCTCCCCCCGCATCCCCGAGCCGGCAGCGTGCGGGGACAGCCCGGGCACGGGGAAGGTGGCACGGGATCGCTTTCCTCTGAACGCTTCTCGCT
GCTCTTTGAGCCTGCAGACACCTGGGGGGATACGGGGAAAAGGCCTCCACGGCCAACTAGCGGGCAGGAAGAGGGCCTATTTCCCATGATTCCTTCATATTTGCATATACGA
TACAAGGCTGTTAGAGAGATAATTAGAATTAATTTGACTGTAAACACAAAGATATTAGTACAAAATACGTGACGTAGAAAGTAATAATTTCTTGGGTAGTTTGCAGTTTTAA
AATTATGTTTTAAAATGGACTATCATATGCTTACCGTAACTTGAAAGTATTTCGATTTCTTGGCTTTATATATCTTGTGGAAAGGACGAAACACCGGAAACCTGGTCAGGGA
GACCGAACGGACTCTAAATCCGTTCAGCCGGGTTCGATTCCCGGGGTTTCCGTTTTTTACTAGCGGGCAGGAAGAGGGCCTATTTCCCATGATTCCTTCATATTTGCATATA
CGATACAAGGCTGTTAGAGAGATAATTAGAATTAATTTGACTGTAAACACAAAGATATTAGTACAAAATACGTGACGTAGAAAGTAATAATTTCTTGGGTAGTTTGCAGTTT
TAAAATTATGTTTTAAAATGGACTATCATATGCTTACCGTAACTTGAAAGTATTTCGATTTCTTGGCTTTATATATCTTGTGGAAAGGACGAAACACCGGAAACCTGGTCAG
GGAGACCGAACGGACTCTAAATCCGTTCAGCCGGGTTCGATTCCCGGGGTTTCCGTTTTTTACTAGCGGGCAGGAAGAGGGCCTATTTCCCATGATTCCTTCATATTTGCAT
ATACGATACAAGGCTGTTAGAGAGATAATTAGAATTAATTTGACTGTAAACACAAAGATATTAGTACAAAATACGTGACGTAGAAAGTAATAATTTCTTGGGTAGTTTGCAG
TTTTAAAATTATGTTTTAAAATGGACTATCATATGCTTACCGTAACTTGAAAGTATTTCGATTTCTTGGCTTTATATATCTTGTGGAAAGGACGAAACACCGGAAACCTGGT
CAGGGAGACCGAACGGACTCTAAATCCGTTCAGCCGGGTTCGATTCCCGGGGTTTCCGTTTTTTACTAGTGCTCTAGAAAGGATCTGCGATCGCTCCGGTGCCCGTCAGTGG
GCAGAGCGCACATCGCCCACAGTCCCCGAGAAGTTGGGGGGAGGGGTCGGCAATTGAACGGGTGCCTAGAGAAGGTGGCGCGGGGTAAACTGGGAAAGTGATGTCGTGTACT
GGCTCCGCCTTTTTCCCGAGGGTGGGGGAGAACCGTATATAAGTGCAGTAGTCGCCGTGAACGTTCTTTTTCGCAACGGGTTTGCCGCCAGAACACAGCTGAAGCTTCGAGG
GGCTCGCATCTCTCCTTCACGCGCCCGCCGCCCTACCTGAGGCCGCCATCCACGCCGGTTGAGTCGCGTTCTGCCGCCTCCCGCCTGTGGTGCCTCCTGAACTGCGTCCGCC
GTCTAGGTAAGTTTAAAGCTCAGGTCGAGACCGGGCCTTTGTCCGGCGCTCCCTTGGAGCCTACCTAGACTCAGCCGGCTCTCCACGCTTTGCCTGACCCTGCTTGCTCAAC
TCTACGTCTTTGTTTCGTTTTCTGTTCTGCGCCGTTACAGATCCAAGCTGTGACCGGCGCCTACTCTAGAGCTAGCCACCATGGACTACAAGGACGACGACGACAAGATGGA
CAAAAAACCGCTGAATACCCTGATCTCTGCTACTGGTCTGTGGATGAGTCGTACCGGAACCATTCATAAAATCAAACACCACGAGGTTAGCCGTTCGAAAATCTATATTGAG
ATGGCGTGTGGCGATCATCTGGTTGTGAACAATAGCCGCTCTTCTCGTACAGCACGTGCACTGCGTCACCACAAATATCGTAAAACCTGTAAACGTTGCCGTGTGTCCGATG
AGGATCTGAACAAATTCCTGACAAAAGCCAATGAGGACCAAACAAGCGTGAAAGTGAAAGTCGTTAGCGCTCCTACCCGTACTAAAAAAGCAATGCCGAAATCCGTTGCTCG
TGCCCCTAAACCACTGGAAAACACTGAAGCAGCACAGGCACAGCCGTCTGGAAGCAAATTCTCTCCGGCCATTCCTGTTTCTACCCAGGAGTCCGTTTCTGTTCCAGCAAGT
GTGAGCACCAGCATTAGCAGTATTAGCACCGGTGCCACCGCTAGCGCCCTGGTTAAAGGCAATACCAATCCGATTACAAGCATGTCTGCCCCGGTTCAAGCATCAGCTCCAG
CACTGACAAAATCCCAAACCGATCGTCTGGAGGTTCTGCTGAATCCGAAAGACGAAATCAGCCTGAATTCCGGCAAACCGTTTCGTGAACTGGAGAGCGAACTGCTGTCACG
TCGTAAAAAAGACCTGCAACAAATCTATGCCGAAGAACGTGAGAACTATCTGGGGAAACTGGAACGTGAAATCACCCGCTTTTTCGTGGATCGTGGCTTTCTGGAGATCAAA
TCCCCGATTCTGATTCCTCTGGAGTATATCGAGCGTATGGGCATCGACAATGATACCGAACTGAGCAAACAAATTTTCCGTGTGGATAAAAACTTCTGTCTGCGCCCTATGC
TAGCACCAAATCTGGCTAACTATCTGCGCAAACTGGACCGTGCCCTGCCTGATCCTATCAAAATCTTCGAGATCGGCCCGTGTTATCGTAAAGAGTCCGACGGTAAAGAACA
TCTGGAGGAGTTTACCATGCTGAACTTTTGCCAAATGGGTTCAGGTTGTACTCGTGAGAACCTGGAAAGCATCATCACCGATTTTCTGAACCACCTGGGCATTGACTTCAAA
ATTGTGGGCGACAGCTGTATGGTGTTTGGCGACACCCTGGATGTCATGCACGGCGACCTGGAACTGTCTAGTGCCGTTGTTGGACCAATTCCGCTGGACCGTGAGTGGGGTA
TCGACAAACCGTGGATCGGAGCAGGATTCGGTCTGGAACGCCTGCTGAAAGTGAAACACGACTTCAAAAACATCAAACGTGCCGCCCGTTCTGAATCGTATTATAACGGGAT
CTCTACGAACCTGTAATAAGGATCCGCGGCCGCGCCCCTCTCCCTCCCCCCCTAACGTTACTGGCCGAAGCCGCTTGGAATAAGGCCGGTGTGCGTTTGTCTATATGTTATT
TTCCACCATATTGCCGTCTTTTGGCAATGTTTATCATCGTGTTTTTCAAAGGAAAACCACGTCCCCGTGGTTCGGGGGGCCTAGACGTTTTTTTAACCTCGACTAAACACAT
GTAAAGCATGTGCACCGAGGCCCCAGATCAGATCCCATACAATGGGGTACCTTCTGGGCATCCTTCAGCCCCTTGTTGAATACGCTTGAGGAGAGCCATTTGACTCTTTCCA
CAACTATCCAACTCACAACGTGGCACTGGGGTTGTGCCGCCTTTGCAGGTGTATCTTATACACGTGGCTTTTGGCCGCAGAGGCACCTGTCGCCAGGTGGGGGGTTCCGCTG
CCTGCAAAGGGTCGCTACAGACGTTGTTTGTCTTCAAGAAGCTTCCAGAGGAACTGCTTCCTTCACGACATTCAACAGACCTTGCATTCCTTTGGCGAGAGGGGAAAGACCC
CTAGGAATGCTCGTCAAGAAGACAGGGCCAGGTTTCCGGGCCCTCTATGGCCACAACCATGGCGTCCGGAATGACCGAGTACAAGCCCACGGTGCGCCTCGCCACCCGCGAC
GACGTCCCCAGGGCCGTACGCACCCTCGCCGCCGCGTTCGCCGACTACCCCGCCACGCGCCACACCGTCGATCCGGACCGCCACATCGAGCGGGTCACCGAGCTGCAAGAAC
TCTTCCTCACGCGCGTCGGGCTCGACATCGGCAAGGTGTGGGTCGCGGACGACGGCGCCGCGGTGGCGGTCTGGACCACGCCGGAGAGCGTCGAAGCGGGGGCGGTGTTCGC
CGAGATCGGCCCGCGCATGGCCGAGTTGAGCGGTTCCCGGCTGGCCGCGCAGCAACAGATGGAAGGCCTCCTGGCGCCGCACCGGCCCAAGGAGCCCGCGTGGTTCCCGGCC
ACCGTCGGCGTCTCGCCCGACCACCAGGGCAAGGGTCTGGGCAGCGCCGTCGTGCTCCCCGGAGTGGAGGCGGCCGAGCGCGCCGGGGTGCCCGCCTTCCTGGAGACCTCCG
CGCCCCGCAACCTCCCCTTCTACGAGCGGCTCGGCTTCACCGTCACCGCCGACGTCGAGGTGCCCGAAGGACCGCGCACCTGGTGCATGACCCGCAAGCCCGGTGCCTGAGT
CGACAATCAACCTCTGGATTACAAAATTTGTGAAAGATTGACTGGTATTCTTAACTATGTTGCTCCTTTTACGCTATGTGGATACGCTGCTTTAATGCCTTTGTATCATGCG
TTAACTAAACTTGTTTATTGCAGCTTATAATGGTTACAAATAAAGCAATAGCATCACAAATTTCACAAATAAAGCATTTTTTTCACTGCATTCTAGTTGTGGTTTGTCCAAA
CTCATCAATGTATCTTATCATGTCTGGAATTGACTCAAATGATGTCAATTAGTCTATCAGAAGCTCATCTGGTCTCCCTTCCGGGGGACAAGACATCCCTGTTTAATATTTA
AACAGCAGTGTTCCCAAACTGGGTTCTTATATCCCTTGCTCTGGTCAACCAGGTTGCAGGGTTTCCTGTCCTCACAGGAACGAAGTCCCTAAAGAAACAGTGGCAGCCAGGT
TTAGCCCCGGAATTGACTGGATTCCTTTTTTAGGGCCCATTGGTATGGCTTTTTCCCCGTATCCCCCCAGGTGTCTGCAGGCTCAAAGAGCAGCGAGAAGCGTTCAGAGGAA
AGCGATCCCGTGCCACCTTCCCCGTGCCCGGGCTGTCCCCGCACGCTGCCGGCTCGGGGATGCGGGGGGAGCGCCGGACCGGAGCGGAGCCCCGGGCGGCTCGCTGCTGCCC
CCTAGCGGGGGAGGGACGTAATTACATCCCTGGGGGCTTTGGGGGGGGGCTGTCCCTGATATCTATAACAAGAAAATATATATATATAATAAGTTATCACGTAAGTAGAACATG
AAATAACAATATAATTATCGTATGAGTTAAATCTTAAAAGTCACGTAAAAGATAATCATGCGTCATTTTGACTCACGCGGTCGTTATAGTTCAAAATCAGTGACACTTACCG
CATTGACAAGCACGCCTCACGGGAGCTCCAAGCGGCGACTGAGATGTCCTAAATGCACAGCGACGGATTCGCGCTATTTAGAAAGAGAGAGCAATATTTCAAGAATGCATGC
GTCAATTTTACGCAGACTATCTTTCTAGGGTTAATCTAGCTGCATCAGGATCATATCGTCGGGTCTTTTTTCCGGCTCAGTCATCGCCCAAGCTGGCGCTATCTGGGCATCG
GGGAGGAAGAAGCCCGTGCCTTTTCCCGCGAGGTTGAAGCGGCATGGAAAGAGTTTGCCGAGGATGACTGCTGCTGCATTGACGTTGAGCGAAAACGCACGTTTACCATGAT
GATTCGGGAAGGTGTGGCCATGCACGCCTTTAACGGTGAACTGTTCGTTCAGGCCACCTGGGATACCAGTTCGTCGCGGCTTTTCCGGACACAGTTCCGGATGGTCAGCCCG
AAGCGCATCAGCAACCCGAACAATACCGGCGACAGCCGGAACTGCCGTGCCGGTGTGCAGATTAATGACAGCGGTGCGGCGCTGGGATATTACGTCAGCGAGGACGGGTATC
CTGGCTGGATGCCGCAGAAATGGACATGGATACCCCGTGAGTTACCCGGCGGGCGCGCTTGGCGTAATCATGGTCATAGCTGTTTCCTGTGTGAAATTGTTATCCGCTCACA
ATTCCACACAACATACGAGCCGGAAGCATAAAGTGTAAAGCCTGGGGTGCCTAATGAGTGAGCTAACTCACATTAATTGCGTTGCGCTCACTGCCCGCTTTCCAGTCGGGAA
ACCTGTCGTGCCAGCTGCATTAATGAATCGGCCAACGCGCGGGGAGAGGCGGTTTGCGTATTGGGCGCTCTTCCGCTTCCTCGCTCACTGACTCGCTGCGCTCGGTCGTTCG
GCTGCGGCGAGCGGTATCAGCTCACTCAAAGGCGGTAATACGGTTATCCACAGAATCAGGGGATAACGCAGGAAAGAACATGTGAGCAAAAGGCCAGCAAAAGGCCAGGAAC
CGTAAAAAGGCCGCGTTGCTGGCGTTTTTCCATAGGCTCCGCCCCCCTGACGAGCATCACAAAAATCGACGCTCAAGTCAGAGGTGGCGAAACCCGACAGGACTATAAAGAT
ACCAGGCGTTTCCCCCTGGAAGCTCCCTCGTGCGCTCTCCTGTTCCGACCCTGCCGCTTACCGGATACCTGTCCGCCTTTCTCCCTTCGGGAAGCGTGGCGCTTTCTCATAG
CTCACGCTGTAGGTATCTCAGTTCGGTGTAGGTCGTTCGCTCCAAGCTGGGCTGTGTGCACGAACCCCCCGTTCAGCCCGACCGCTGCGCCTTATCCGGTAACTATCGTCTT
GAGTCCAACCCGGTAAGACACGACTTATCGCCACTGGCAGCAGCCACTGGTAACAGGATTAGCAGAGCGAGGTATGTAGGCGGTGCTACAGAGTTCTTGAAGTGGTGGCCTA
ACTACGGCTACACTAGAAGAACAGTATTTGGTATCTGCGCTCTGCTGAAGCCAGTTACCTTCGGAAAAAGAGTTGGTAGCTCTTGATCCGGCAAACAAACCACCGCTGGTAG
CGGTGGTTTTTTTGTTTGCAAGCAGCAGATTACGCGCAGAAAAAAAGGATCTCAAGAAGATCCTTTGATCTTTTCTACGGGGTCTGACGCTCAGTGGAACGAAAACTCACGT
TAAGGGATTTTGGTCATGAGATTATCAAAAAGGATCTTCACCTAGATCCTTTTAAATTAAAAATGAAGTTTTAAATCAATCTAAAGTATATATGAGTAAACTTGGTCTGACA
GTTACCAATGCTTAATCAGTGAGGCACCTATCTCAGCGATCTGTCTATTTCGTTCATCCATAGTTGCCTGACTCCCCGTCGTGTAGATAACTACGATACGGGAGGGCTTACC
ATCTGGCCCCAGTGCTGCAATGATACCGCGAGACCCACGCTCACCGGCTCCAGATTTATCAGCAATAAACCAGCCAGCCGGAAGGGCCGAGCGCAGAAGTGGTCCTGCAACT
TTATCCGCCTCCATCCAGTCTATTAATTGTTGCCGGGAAGCTAGAGTAAGTAGTTCGCCAGTTAATAGTTTGCGCAACGTTGTTGCCATTGCTACAGGCATCGTGGTGTCAC
GCTCGTCGTTTGGTATGGCTTCATTCAGCTCCGGTTCCCAACGATCAAGGCGAGTTACATGATCCCCCATGTTGTGCAAAAAAGCGGTTAGCTCCTTCGGTCCTCCGATCGT
TGTCAGAAGTAAGTTGGCCGCAGTGTTATCACTCATGGTTATGGCAGCACTGCATAATTCTCTTACTGTCATGCCATCCGTAAGATGCTTTTCTGTGACTGGTGAGTACTCA

ACCAAGTCATTCTGAGAATAGTGTATGCGGCGACCGAGTTGCTCTTGCCCGGCGTCAATACGGGATAATACCGCGCCACATAGCAGAACTTTAAAAGTGCTCATCATTGGAA
AACGTTCTTCGGGGCGAAAACTCTCAAGGATCTTACCGCTGTTGAGATCCAGTTCGATGTAACCCACTCGTGCACCCAACTGATCTTCAGCATCTTTTACTTTCACCAGCGT
TTCTGGGTGAGCAAAAACAGGAAGGCAAAATGCCGCAAAAAAGGGAATAAGGGCGACACGGAAATGTTGAATACTCATACTCTTCCTTTTTCAATATTATTGAAGCATTTAT
CAGGGTTATTGTCTCATGAGCGGATACATATTTGAATGTATTTAGAAAAATAAACAAATAGGGGTTCCGCGCACATTTCCCCGAAAAGTGCCACCTAAATTGTAAGCGTTAA
TATTTTGTTAAAATTCGCGTTAAATTTTTGTTAAATCAGCTCATTTTTTAACCAATAGGCCGAAATCGGCAAAATCCCTTATAAATCAAAAGAATAGACCGAGATAGGGTTG
AGTGTTGTTCCAGTTTGGAACAAGAGTCCACTATTAAAGAACGTGGACTCCAACGTCAAAGGGCGAAAAACCGTCTATCAGGGCGATGGCCCACTACGTGAACCATCACCCT
AATCAAGTTTTTTGGGGTCGAGGTGCCGTAAAGCACTAAATCGGAACCCTAAAGGGAGCCCCCGATTTAGAGCTTGACGGGGAAAGCCGGCGAACGTGGCGAGAAAGGAAGG
GAAGAAAGCGAAAGGAGCGGGCGCTAGGGCGCTGGCAAGTGTAGCGGTCACGCTGCGCGTAACCACCACACCCGCCGCGCTTAATGCGCCGCTACAGGGCGCGTCCCATTCG
CCATTCAGGCTGCGCAACTGTTGGGAAGGGCGATCGGTGCGGGCCTCTTCGCTATTACGCCAGCTGGCGAAAGGGGGATGTGCTGCAAGGCGATTAAGTTGGGTAACGCCA
GGGTTTTCCCAGTCACGACGTTGTAAAACGACGGCCAGTGAGCGCGCCTCGTTCATTCACGTTTTTGAACCCGTGGAGGACGGGCAGACTCGCGGTGCAAATGTGTTTTACA
GCGTGATGGAGCAGATGAAGATGCTCGACACGCTGCA

AAV-M15 transfer vector:

AAV2 ITR CMV enhancer CMV promoter Flag-PylRS U6 tRNA-M15

CTTTTGCTGGCCTTTTGCTCACATGTCCTGCAGGCAGCTGCGCGCTCGCTCGCTCACTGAGGCCGCCCGGGCGTCGGGCGACCTTTGGTCGCCCGGCCTCAGTGAGCGAGCG
AGCGCGCAGAGAGGGAGTGGCCAACTCCATCACTAGGGGTTCCTGCGGCCTCTAGACTCGAGGCGTTGACATTGATTATTGACTAGTTATTAATAGTAATCAATTACGGGGT
CATTAGTTCATAGCCCATATATGGAGTTCCGCGTTACATAACTTACGGTAAATGGCCCGCCTGGCTGACCGCCCAACGACCCCCGCCCATTGACGTCAATAATGACGTATGT
TCCCATAGTAACGCCAATAGGGACTTTCCATTGACGTCAATGGGTGGAGTATTTACGGTAAACTGCCCACTTGGCAGTACATCAAGTGTATCATATGCCAAGTACGCCCCCT
ATTGACGTCAATGACGGTAAATGGCCCGCCTGGCATTATGCCCAGTACATGACCTTATGGGACTTTCCTACTTGGCAGTACATCTACGTATTAGTCATCGCTATTACCATGG
TGATGCGGTTTTGGCAGTACATCAATGGGCGTGGATAGCGGTTTGACTCACGGGGATTTCCAAGTCTCCACCCCATTGACGTCAATGGGAGTTTGTTTTGGCACCAAAATCA
ACGGGACTTTCCAAAATGTCGTAACAACTCCGCCCCATTGACGCAAATGGGCGGTAGGCGTGTACGGTGGGAGGTCTATATAAGCAGAGCTCTCTGGCTAACTACCGGTGCC
ACCATGGACTACAAGGACGACGACGACAAGATGGACAAAAAACCGCTGAATACCCTGATCTCTGCTACTGGTCTGTGGATGAGTCGTACCGGAACCATTCATAAAATCAAAC
ACCACGAGGTTAGCCGTTCGAAAATCTATATTGAGATGGCGTGTGGCGATCATCTGGTTGTGAACAATAGCCGCTCTTCTCGTACAGCACGTGCACTGCGTCACCACAAATA
TCGTAAAACCTGTAAACGTTGCCGTGTGTCCGATGAGGATCTGAACAAATTCCTGACAAAAGCCAATGAGGACCAAACAAGCGTGAAAGTGAAAGTCGTTAGCGCTCCTACC
CGTACTAAAAAAGCAATGCCGAAATCCGTTGCTCGTGCCCCTAAACCACTGGAAAACACTGAAGCAGCACAGGCACAGCCGTCTGGAAGCAAATTCTCTCCGGCCATTCCTG
TTTCTACCCAGGAGTCCGTTTCTGTTCCAGCAAGTGTGAGCACCAGCATTAGCAGTATTAGCACCGGTGCCACCGCTAGCGCCCTGGTTAAAGGCAATACCAATCCGATTAC
AAGCATGTCTGCCCCGGTTCAAGCATCAGCTCCAGCACTGACAAAATCCCAAACCGATCGTCTGGAGGTTCTGCTGAATCCGAAAGACGAAATCAGCCTGAATTCCGGCAAA
CCGTTTCGTGAACTGGAGAGCGAACTGCTGTCACGTCGTAAAAAAGACCTGCAACAAATCTATGCCGAAGAACGTGAGAACTATCTGGGGAAACTGGAACGTGAAATCACCC
GCTTTTTCGTGGATCGTGGCTTTCTGGAGATCAAATCCCCGATTCTGATTCCTCTGGAGTATATCGAGCGTATGGGCATCGACAATGATACCGAACTGAGCAAACAAATTTT
CCGTGTGGATAAAAACTTCTGTCTGCGCCCTATGCTAGCACCAAATCTGGCTAACTATCTGCGCAAACTGGACCGTGCCCTGCCTGATCCTATCAAAATCTTCGAGATCGGC
CCGTGTTATCGTAAAGAGTCCGACGGTAAAGAACATCTGGAGGAGTTTACCATGCTGAACTTTTGCCAAATGGGTTCAGGTTGTACTCGTGAGAACCTGGAAAGCATCATCA
CCGATTTTCTGAACCACCTGGGCATTGACTTCAAAATTGTGGGCGACAGCTGTATGGTGTTTGGCGACACCCTGGATGTCATGCACGGCGACCTGGAACTGTCTAGTGCCGT
TGTTGGACCAATTCCGCTGGACCGTGAGTGGGGTATCGACAAACCGTGGATCGGAGCAGGATTCGGTCTGGAACGCCTGCTGAAAGTGAAACACGACTTCAAAAACATCAAA
CGTGCCGCCCGTTCTGAATCGTATTATAACGGGATCTCTACGAACCTGTAAAGCGGCCGCTCGATGCTTTATTTGTGAAATTTGTGATGCTATTGCTTTATTTGTAACCATT
ATAAGCTGCAATAAACAAGTTAACAACAACAATTGCATTCATTTTATGTTTCAGGTTCAGGGGGAGGTGTGGGAGGTTTTTTAAGGTACCTATGCTAGCCGCGGCCTCCACG
GCCAACTAGCGGGCAGGAAGAGGGCCTATTTCCCATGATTCCTTCATATTTGCATATACGATACAAGGCTGTTAGAGAGATAATTAGAATTAATTTGACTGTAAACACAAAG
ATATTAGTACAAAATACGTGACGTAGAAAGTAATAATTTCTTGGGTAGTTTGCAGTTTTAAAATTATGTTTTAAAATGGACTATCATATGCTTACCGTAACTTGAAAGTATT
TCGATTTCTTGGCTTTATATATCTTGTGGAAAGGACGAAACACCGGAAACCTGGTCAGGGAGACCGAACGGACTCTAAATCCGTTCAGCCGGGTTCGATTCCCGGGGTTTCC
GTTTTTTACTAGCGGGCAGGAAGAGGGCCTATTTCCCATGATTCCTTCATATTTGCATATACGATACAAGGCTGTTAGAGAGATAATTAGAATTAATTTGACTGTAAACACA
AAGATATTAGTACAAAATACGTGACGTAGAAAGTAATAATTTCTTGGGTAGTTTGCAGTTTTAAAATTATGTTTTAAAATGGACTATCATATGCTTACCGTAACTTGAAAGT
ATTTCGATTTCTTGGCTTTATATATCTTGTGGAAAGGACGAAACACCGGAAACCTGGTCAGGGAGACCGAACGGACTCTAAATCCGTTCAGCCGGGTTCGATTCCCGGGGTT
TCCGTTTTTTACTAGCGGGCAGGAAGAGGGCCTATTTCCCATGATTCCTTCATATTTGCATATACGATACAAGGCTGTTAGAGAGATAATTAGAATTAATTTGACTGTAAAC
ACAAAGATATTAGTACAAAATACGTGACGTAGAAAGTAATAATTTCTTGGGTAGTTTGCAGTTTTAAAATTATGTTTTAAAATGGACTATCATATGCTTACCGTAACTTGAA
AGTATTTCGATTTCTTGGCTTTATATATCTTGTGGAAAGGACGAAACACCGGAAACCTGGTCAGGGAGACCGAACGGACTCTAAATCCGTTCAGCCGGGTTCGATTCCCGGG
GTTTCCGTTTTTTACTAGTTATGCTAGCCGCGCGGCCGCAGGAACCCCTAGTGATGGAGTTGGCCACTCCCTCTCTGCGCGCTCGCTCGCTCACTGAGGCCGGGCGACCAAA
GGTCGCCCGACGCCCGGGCTTTGCCCGGGCGGCCTCAGTGAGCGAGCGAGCGCGCAGCTGCCTGCAGGGGCGCCTGATGCGGTATTTTCTCCTTACGCATCTGTGCGGTATT
TCACACCGCATACGTCAAAGCAACCATAGTACGCGCCCTGTAGCGGCGCATTAAGCGCGGCGGGTGTGGTGGTTACGCGCAGCGTGACCGCTACACTTGCCAGCGCCTTAGC
GCCCGCTCCTTTCGCTTTCTTCCCTTCCTTTCTCGCCACGTTCGCCGGCTTTCCCCGTCAAGCTCTAAATCGGGGGCTCCCTTTAGGGTTCCGATTTAGTGCTTTACGGCAC
CTCGACCCCAAAAAACTTGATTTGGGTGATGGTTCACGTAGTGGGCCATCGCCCTGATAGACGGTTTTTCGCCCTTTGACGTTGGAGTCCACGTTCTTTAATAGTGGACTCT
TGTTCCAAACTGGAACAACACTCAACTCTATCTCGGGCTATTCTTTTGATTTATAAGGGATTTTGCCGATTTCGGTCTATTGGTTAAAAAATGAGCTGATTTAACAAAAATT
TAACGCGAATTTTAACAAAATATTAACGTTTACAATTTTATGGTGCACTCTCAGTACAATCTGCTCTGATGCCGCATAGTTAAGCCAGCCCCGACACCCGCCAACACCCGCT
GACGCGCCCTGACGGGCTTGTCTGCTCCCGGCATCCGCTTACAGACAAGCTGTGACCGTCTCCGGGAGCTGCATGTGTCAGAGGTTTTCACCGTCATCACCGAAACGCGCGA
GACGAAAGGGCCTCGTGATACGCCTATTTTTATAGGTTAATGTCATGATAATAATGGTTTCTTAGACGTCAGGTGGCACTTTTCGGGGAAATGTGCGCGGAACCCCTATTTG
TTTATTTTTCTAAATACATTCAAATATGTATCCGCTCATGAGACAATAACCCTGATAAATGCTTCAATAATATTGAAAAAGGAAGAGTATGAGTATTCAACATTTCCGTGTC
GCCCTTATTCCCTTTTTTGCGGCATTTTGCCTTCCTGTTTTTGCTCACCCAGAAACGCTGGTGAAAGTAAAAGATGCTGAAGATCAGTTGGGTGCACGAGTGGGTTACATCG
AACTGGATCTCAACAGCGGTAAGATCCTTGAGAGTTTTCGCCCCGAAGAACGTTTTCCAATGATGAGCACTTTTAAAGTTCTGCTATGTGGCGCGGTATTATCCCGTATTGA
CGCCGGGCAAGAGCAACTCGGTCGCCGCATACACTATTCTCAGAATGACTTGGTTGAGTACTCACCAGTCACAGAAAAGCATCTTACGGATGGCATGACAGTAAGAGAATTA
TGCAGTGCTGCCATAACCATGAGTGATAACACTGCGGCCAACTTACTTCTGACAACGATCGGAGGACCGAAGGAGCTAACCGCTTTTTTGCACAACATGGGGGATCATGTAA
CTCGCCTTGATCGTTGGGAACCGGAGCTGAATGAAGCCATACCAAACGACGAGCGTGACACCACGATGCCTGTAGCAATGGCAACAACGTTGCGCAAACTATTAACTGGCGA
ACTACTTACTCTAGCTTCCCGGCAACAATTAATAGACTGGATGGAGGCGGATAAAGTTGCAGGACCACTTCTGCGCTCGGCCCTTCCGGCTGGCTGGTTTATTGCTGATAAA
TCTGGAGCCGGTGAGCGTGGAAGCCGCGGTATCATTGCAGCACTGGGGCCAGATGGTAAGCCCTCCCGTATCGTAGTTATCTACACGACGGGGAGTCAGGCAACTATGGATG
AACGAAATAGACAGATCGCTGAGATAGGTGCCTCACTGATTAAGCATTGGTAACTGTCAGACCAAGTTTACTCATATATACTTTAGATTGATTTAAAACTTCATTTTTAATT
TAAAAGGATCTAGGTGAAGATCCTTTTTGATAATCTCATGACCAAAATCCCTTAACGTGAGTTTTCGTTCCACTGAGCGTCAGACCCCGTAGAAAAGATCAAAGGATCTTCT
TGAGATCCTTTTTTTCTGCGCGTAATCTGCTGCTTGCAAACAAAAAAACCACCGCTACCAGCGGTGGTTTGTTTGCCGGATCAAGAGCTACCAACTCTTTTTCCGAAGGTAA
CTGGCTTCAGCAGAGCGCAGATACCAAATACTGTTCTTCTAGTGTAGCCGTAGTTAGGCCACCACTTCAAGAACTCTGTAGCACCGCCTACATACCTCGCTCTGCTAATCCT
GTTACCAGTGGCTGCTGCCAGTGGCGATAAGTCGTGTCTTACCGGGTTGGACTCAAGACGATAGTTACCGGATAAGGCGCAGCGGTCGGGCTGAACGGGGGGTTCGTGCACA
CAGCCCAGCTTGGAGCGAACGACCTACACCGAACTGAGATACCTACAGCGTGAGCTATGAGAAAGCGCCACGCTTCCCGAAGGGAGAAAGGCGGACAGGTATCCGGTAAGCG
GCAGGGTCGGAACAGGAGAGCGCACGAGGGAGCTTCCAGGGGGAAACGCCTGGTATCTTTATAGTCCTGTCGGGTTTCGCCACCTCTGACTTGAGCGTCGATTTTTGTGATG
CTCGTCAGGGGGGCGGAGCCTATGGAAAAACGCCAGCAACGCGGCCTTTTTACGGTTCCTGGC
